# Phomoarcherin B as a novel HIV-1 reverse transcriptase RNase H activity inhibitor: conclusions from comprehensive computational analysis

**DOI:** 10.1101/2021.09.09.459559

**Authors:** Naeem Abdul Ghafoor, Ömür Baysal, Barış Ethem Süzek, Ragıp Soner Silme

## Abstract

The HIV epidemic has claimed more than 32.7 million live since its emergence in 1981, while many ART and HAART therapies are available and provide relief and control for patients, most of these therapeutics come with long-term side effects, resistance, socio-economical barriers, and other obstacles. In this study, genomic analysis was performed on 98 HIV-1 genomes to determine the most coherent target that could be utilized to restrict and cease the viral replication, the reverse transcriptase enzyme. Following the identification of the target protein, the RNase H activity of the reverse transcriptase was nominated as the potent target given the limited research associated with it. A library of 94 thousand small molecule inhibitors was generated and virtual screening was performed to identify hits, based on the reproducibility of the screening results, 4 compounds with the best scores were considered and their interaction within the active site was analyzed. Subsequently, all-atom molecular dynamics simulations and MM-PBSA was performed to validate the stability and binding free energy of the hits within the RNase H active. In silico ADMET assays were performed on the hit compounds to analyze their drug-likeness, physicochemical and pharmacological properties. Phomoarcherin B, a pentacyclic aromatic sesquiterpene naturally found in the endophytic fungus *Phomopsis archeri*, known for its anticancer properties scored the best in all the experiments and was nominated as a potential inhibitor of the HIV-1 reverse transcriptase RNase H activity.

**Author Summary:** The Human immunodeficiency virus 1 (HIV-1) has remained a global public health issue for the past 4 decades with millions of patients around the globe affected. Long years of research in the field of anti-viral therapies had introduced several drugs to combat the viral infection, however, most of these drugs come with their shortcomings. In this study, computational drug discovery approaches were utilized to research novel drugs that could be used as anti-viral agents against the former virus, the results had revealed Phomoarcherin B, a natural compound from an endophytic fungus to hold promising therapeutic potentials.

## 1. Introduction

The Human immunodeficiency virus 1 (HIV-1) remains a global public health issue ever since its first identification in 1981, its the caustic agent for the Acquired immunodeficiency syndrome (AIDS) which compromises the immune system of the host and eventually leave them defenseless against secondary infection [1]. The World Health Organization refers to HIV-1 as a “global epidemic” [2], and according to the Joint United Nations Programme on HIV/AIDS (UNAIDS) global statistics around 24.8 million to 42.2 million individuals have died due to AIDS-related illnesses since its emergence and by 2019 a total of 31.6 million to 44.5 million individuals still live with the virus [3], most of them living in sub-Saharan Africa [4].

HIV is a complex retrovirus, like other retroviruses it stores its genome as a pair of ssRNA molecules of around 9kb long. The genome contains the gag gene which encodes the structural proteins, mainly the protein capsid, the matrix protein, and the nucleocapsid, the genome also contains the pol gene which encodes the reverse transcriptase enzyme (RT), the protease enzyme, and the integrase enzyme and also the env gene which encodes the membrane glycoprotein 120 and glycoprotein 41. HIV genome also encodes 6 regulatory proteins such as tat, rev, nef, vif. vpr, and vpu which are responsible for its pathogenicity and replication in the host [5]. The viral particle is around 100nm in diameter and its outer membrane is a lipid-rich lipoprotein membrane, the glycoprotein120 trimer is present in the external surface of the membrane by forming a heterodimer with the glycoprotein41 which is embedded into the transmembranal domain, the matrix proteins are anchored into the inner domain of the lipoprotein membrane. Inside the lipoprotein membrane and the matrix anchor, the viral capsid which holds the viral gRNA pair along with its regulatory proteins exists, reverse transcriptase (RT) and integrase (IN) enzyme, the protease enzyme is however packed outside the capsid. The HIV replication cycle begins with the binding of the glycoprotein120 to the host’s CD4+ receptor which presents on the surface of activated T helper cells, however, recent researches had shown that while T helper cells are the primary target for HIV, other immune cells such as macrophages, immature dendritic cells, and other resting T cell subsets could be infected as well, once the glycoprotein120 binds to its target, the viral envelope undergoes conformational changes which exposes specific domains in the glycoprotein120 which then binds to the co-receptor CCR5 or CXCR4 (depending on the virus’ tropism). The glycoprotein120’s binding to receptor and co-receptor initiates the N-terminal fusion peptide of glycoprotein41 to penetrate the host cell membrane and hold both the membranes close to each other to initiate the fusion of the membranes which leads to the insertion of the viral capsid into the host cell. Once inside the host, the capsid disintegrates and the viral RNAs are reverse transcribed into DNA molecules via RT which starts with an RNA/DNA hybrid and further cleavage of the RNA, and synthesis of dsDNA takes place. The proviral dsDNA is further integrated into the host genome via the integrase enzyme where it remains as a reservoir for the virus until the cell is activated upon which the cellular transcription and translation mechanisms are hijacked to produce the viral proteins and viral RNA genomes [6–8].

The first clinical attempts to treat the virus started in 1987 and later that year zidovudine (ZDV, also referred to as AZT or azidothymidine) was approved for clinical usage which marked the beginning of the development of antiretroviral therapies (ARTs) strategies [9]. Nowadays, 6 classes of ART drugs targetting 5 different phases of the HIV life cycle has been developed and they’re classified as nucleoside and nucleotide reverse transcriptase inhibitors (NRTIs), non-nucleoside reverse transcription inhibitors (NNRTIs), protease inhibitors (PIs), entry or fusion inhibitors, and integrase strand transfer inhibitors (INSTIs) [10]. A list of all the FDA-approved ART drugs against HIV-1 is shown in Table 1.

**Table 1.**
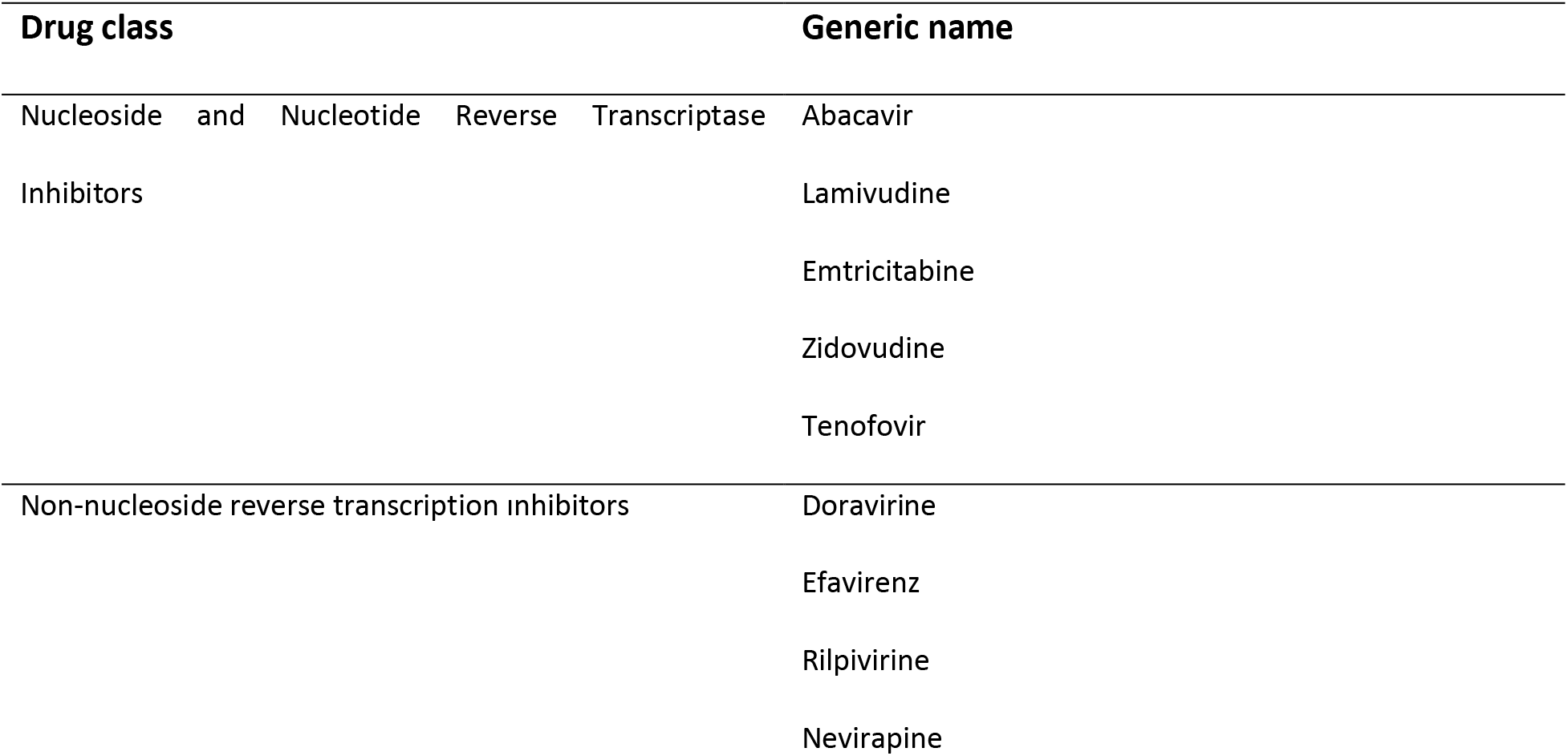
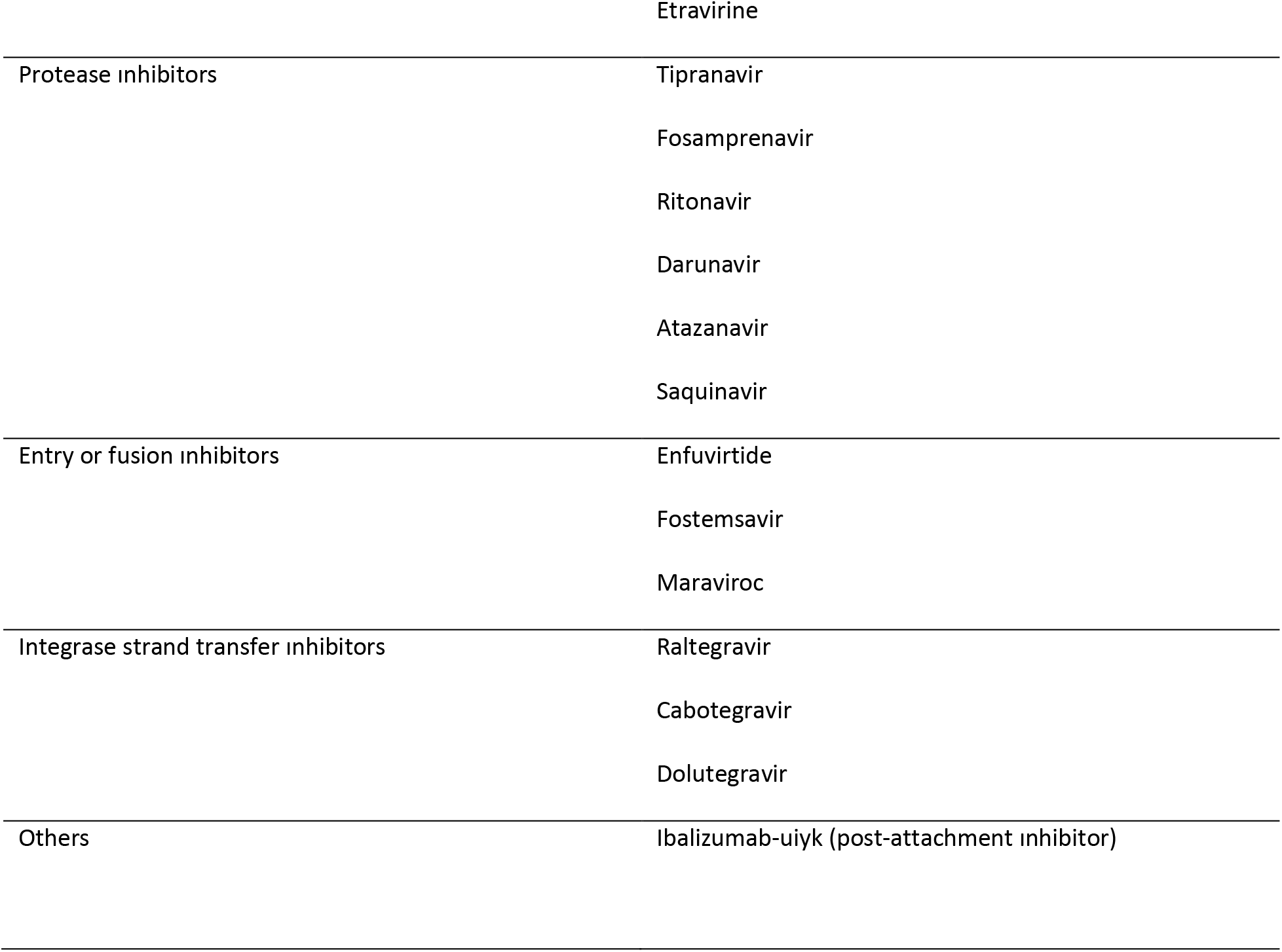
FDA approved ART drugs against HIV, data based on U.S. FDA website.

| Drug class |  |  |  |  | Generic name |
| --- | --- | --- | --- | --- | --- |
| Nucleoside and Nucleotide Reverse Transcriptase Inhibitors |  |  |  |  | Abacavir |
|  |  |  |  |  | Lamivudine |
|  |  |  |  |  | Emtricitabine |
|  |  |  |  |  | Zidovudine |
|  |  |  |  |  | Tenofovir |
| Non-nucleoside reverse transcription inhibitors |  |  |  |  | Doravirine |
|  |  |  |  |  | Efavirenz |
|  |  |  |  |  | Rilpivirine |
|  |  |  |  |  | Nevirapine |

|  |  |
| --- | --- |
|  | Etravirine |
| Protease inhibitors | Tipranavir<br>Fosamprenavir<br>Ritonavir<br>Darunavir<br>Atazanavir<br>Saquinavir |
| Entry or fusion inhibitors | Enfuvirtide<br>Fostemsavir<br>Maraviroc |
| Integrase strand transfer inhibitors | Raltegravir<br>Cabotegravir<br>Dolutegravir |
| Others | Ibalizumab-uiyk (post-attachment inhibitor) |

Another substantial discovery in the field of ART for HIV treatment started with the high active antiretroviral therapy (HAART) era in the mid-1990s. The replication mechanism of HIV-1 is far from perfect, mainly due to its error-prone reverse transcriptase which randomly introduces mutations into the viral genome which could over time lead to resistance against drugs and HAART provided the solution for the issues. In HAART, a cocktail of several ART drugs from different drugs are prescribed at the same time, since the probability of developing mutations to become resistant to several drugs at the same time is far less likely than developing mutations against a single drug, HAART has minimized the drug resistance issue that single ART drugs caused substantially [1,11,12]. Despite the hope, HAART holds for its patients and the opportunities it provides, its yet far from perfect as the current drugs prescribed under HAART comes with many short and/or long term side effects such as hepatotoxicity, cardiovascular complications, distal sensory peripheral neuropathy, lipodystrophy, reduction in the bone mineral density, mitochondrial toxicity, alteration of glucose and serum lipids levels, and the probability of developing resistance remains as well [13–17]. Besides the health issues, access to ARTs and HIV healthcare is bound to socio-economical challenges as it can cost the patients around $21,376 annually in the US, an average of $2,381 for African countries, and for European countries, it varies between $11,618 and $22,673 depending on the country but regardless of the geography, HIV care is a burden both for the health system and families of the patients and for patients in low-income countries, the burden doesn’t get any lighter [18].

With the advances in sequencing technologies in recent years, a huge amount of data regarding different aspects of the viral replication cycle has been accumulated over the years and with the development of many different bioinformatic tools, designing and testing hypotheses has become feasible for larger audiences. The Los Alamos HIV database alone hosts around 13,688 complete genome sequences and over the years many bioinformatic tools have been used for subtyping and classification of the virus [19–21]. Computational methods have also been used widely for designing several neutralizing antibodies against the virus [22–25].

Among the ART drugs and regimens developed for HIV-1, RT has been a critical target, the very first drug Azidothymidine is an NRTIs, the basic mechanism of NRTIs is in its binding to the newly synthesized viral DNA by RT and causing chain termination (mimicking nucleosides/nucleotides) thereby stoping further synthesis of viral DNA [26,27]. NNRTIs perform the same task of inhibiting the RT action but slightly in a different manner, rather than causing chain terminations, NNRTIs interact with catalytic sites or allosteric sites of RT thereby inhibiting its activity and/or blocking it from contacting the substrate i.e. the viral RNA [28]. The HIV-1 RT has 2 main functions performed by 2 separate domains, the DNA polymerase domain (both RNA-dependent and DNA-dependent) and the RNase H domain. The DNA polymerase synthesizes the proviral minus-strand DNA from the viral plus-strand RNA hence producing a DNA/RNA duplex in the process, the primer for the synthesis of this DNA is the host’s tRNA. The RNase H domain utilizes its ribonuclease activity to cleave the RNA from the DNA/RNA duplex and the synthesis of plus-strand DNA is initiated and a double-stranded DNA of the virus is produced which is further translocated into the nucleus and integrated into the host genome [29].

In total, 5 FDA-approved NNRTIs are commercially available (Table 1), doravirine is a pyridinone NNRTI that binds to the hydrophobic pocket of HIV-1’s RT DNA polymerase domains, efavirenz acts similarly by inhibiting the RNA-dependent and DNA-dependent activity, rilpivirine as well inhibits the HIV-1 RT by binding non-competitively near the DNA polymerases’ hydrophobic pocket thereby changing the enzyme’s conformation, nevirapine also inhibits the HIV-1 by binding at the NNRTI-binding pocket and etravirine binds to the reverse transcriptase in various conformations at the NNRTI-binding pocket [35–39]. Considering the mechanism of action for the currently available NNRTIs, all of them target the polymerase activity of the HIV-1 RT and none of them target the RNase H activity/catalytic site, therefore, the discovery and development of novel affordable therapeutic agents targetting the HIV-1 RT’s RNase H is a rather less researched area and holds a promising option for patients running out of options for the currently available drugs, patients sensitive to some or many of the currently available drugs, patients with multi-drug resistance and patients unable to afford or access the currently available drugs.

In early studies, several attempts have been made to target the RNase H activity of the HIV-1 in vitro, however, most of the lead compounds in these studies lacked the specificity to the viral RNase H domain and hence were dropped from further drug research programs [40–43]. More recent studies focused on the development of small-molecule RNA fragment inhibitors that competitively binds to the RNase H domain that produced promising results in *in vitro* experiments by inhibiting the HIV-1 replication cycle, however with the duration of the RNA molecules’ stability, such attempts didn’t achieve much success in the following trials [44]. Some other studies explored inhibitors based on metal-ion co-factor inhibition such as 4-chlorophenylhydrazone of mesoxalic acid, tropolone and its derivatives, h-Hydroxyimides, and diketo acids that function by blocking the access or recruitment of the metal ion to the active site of RNase H, however, none of the potential compounds have been extensively researched or earned FDA approval [45–49].

Computational drug discovery has gained huge momentum in recent years, especially with the availability of supercomputers and the integration of graphic cards to accelerate computation processes which made many bioinformatics and computational research highly spontaneous and less time-intensive, among the fields that benefitted the most from these developments was virtual screening which demanded an immense amount of time for screening large libraries to determine potential novel leads, and molecular dynamics which is exceedingly computationally expensive but holds a great promise in providing atom-level insights for molecular interactions under controlled/designed conditions, the developments of such multiscale models for simulating complex chemical systems has also won the 2013 Nobel prize in chemistry for the revolution it brought in understanding biomolecular systems [50–52].

Virtual screening to identify lead compounds that potentially inhibit several HIV-1 proteins and enzymes had been previously researched, including against the RNase domain, however, such researches were based solely on molecular docking, pharmacophore, and ADMET assays which considers mainly the best docking pose the ligands can have against the target protein in a vacuum-like condition and are trained on limited training datasets, which may or may not reflect their interaction in the physiological cell condition [53–55]. Extensive computational analysis and molecular dynamics for leads against potential HIV-1 RT RNase domain were quite recently performed by Zhang et al. (2016), however, the experiment was limited only to 77 α-hydroxytropolone derivatives, which limited the efforts of discovering novel small molecule inhibitors, hence, no large-scale extensive in silico analysis with all-atom molecular dynamics simulations, and/or some form in silico binding free energy calculation validating the potential lead compounds have been researched against the HIV-1 RT RNase domain [56].

In the current study, genomic data analysis was utilized to indicate the rationale of targeting RT for the drug discovery and development efforts for anti-HIV-1, following the establishment of the RT enzyme as the best candidate for drug targeting, a dataset of around 94 thousand small drug-like molecules was obtained from the ZINC15 database, structure-based virtual screen against the RT RNase domain was performed via molecular docking, the interaction and dynamics of the top lead molecules were further validated using molecular dynamics. The Molecular Mechanics Poisson–Boltzmann Surface Area (MM/PBSA) end-point binding free energy calculation method which combines molecular mechanics and continuum solvents was deployed to estimate the binding energies of the top leads, their physiochemical and pharmacological properties were analyzed via in silico absorption, distribution, metabolism, excretion, and toxicity (ADMET) assays, and their interactions within the RNase H active site was investigated.

## 2. Results

### 2.1. Whole-genome alignment & BLASTx

The results from the whole-genome alignment are provided in Fig 3, the alignment is visualized such that each LCB is clustered together, the alignment is zoomed out so as each clustered LCB is shown as a black continuous bar, a region with a length of around 2.4 kb within the ≈2.8-5.3 kb range from the consensus sequence is highly conserved with almost no gaps in most of the sequences. The alignment for this conserved region further shows a very high degree of conservation within this region throughout all of the HIV-1 genomes aligned with a gap only in one of the sequences (Fig 4).

**Fig 1.**
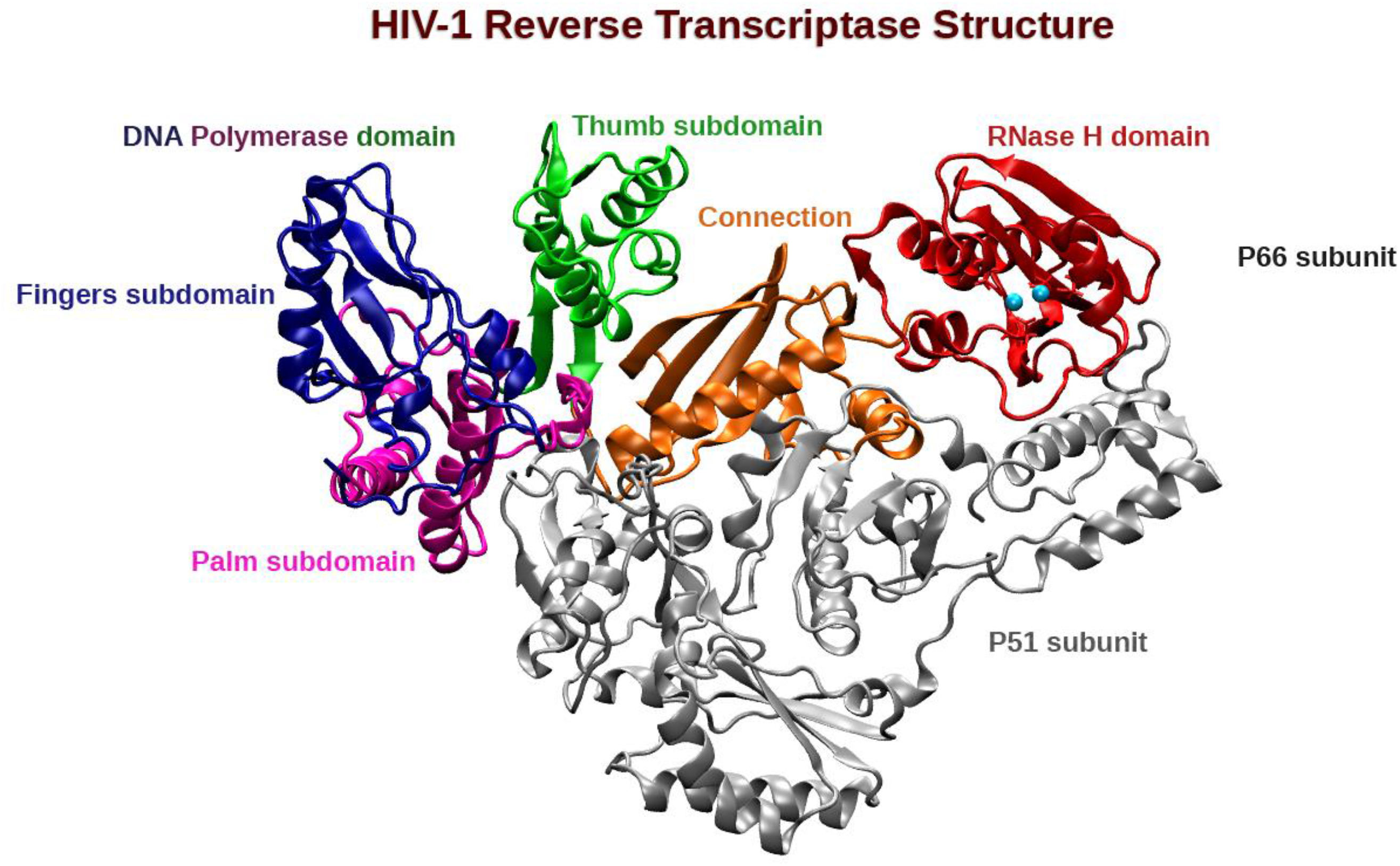
HIV-1 RT enzyme in cartoon representation. Chain B (silver) also called the P51 subunit contains no catalytic sites whereas chain A, also known as the p66 subunit, contains both the polymerase and the ribonuclease domains, the DNA polymerase domain consists of 3 subdomains, the fingers subdomain (blue), the palm subdomain (pink), the thumb subdomain (green), a connecting region (orange) bridges the DNA polymerase domain to the RNase H domain (red) which also contains 2 Mn atoms as cofactor (cyan beads, combinations of Mg^2+^ and Mn^2+^ or only Mg^2+^ has been experimentally identified and produces equivalent activity). The was image generated with VMD (v1.9.3) and secondary structures assigned via STRIDE (v1.0) [30–34].

**Fig 2.**
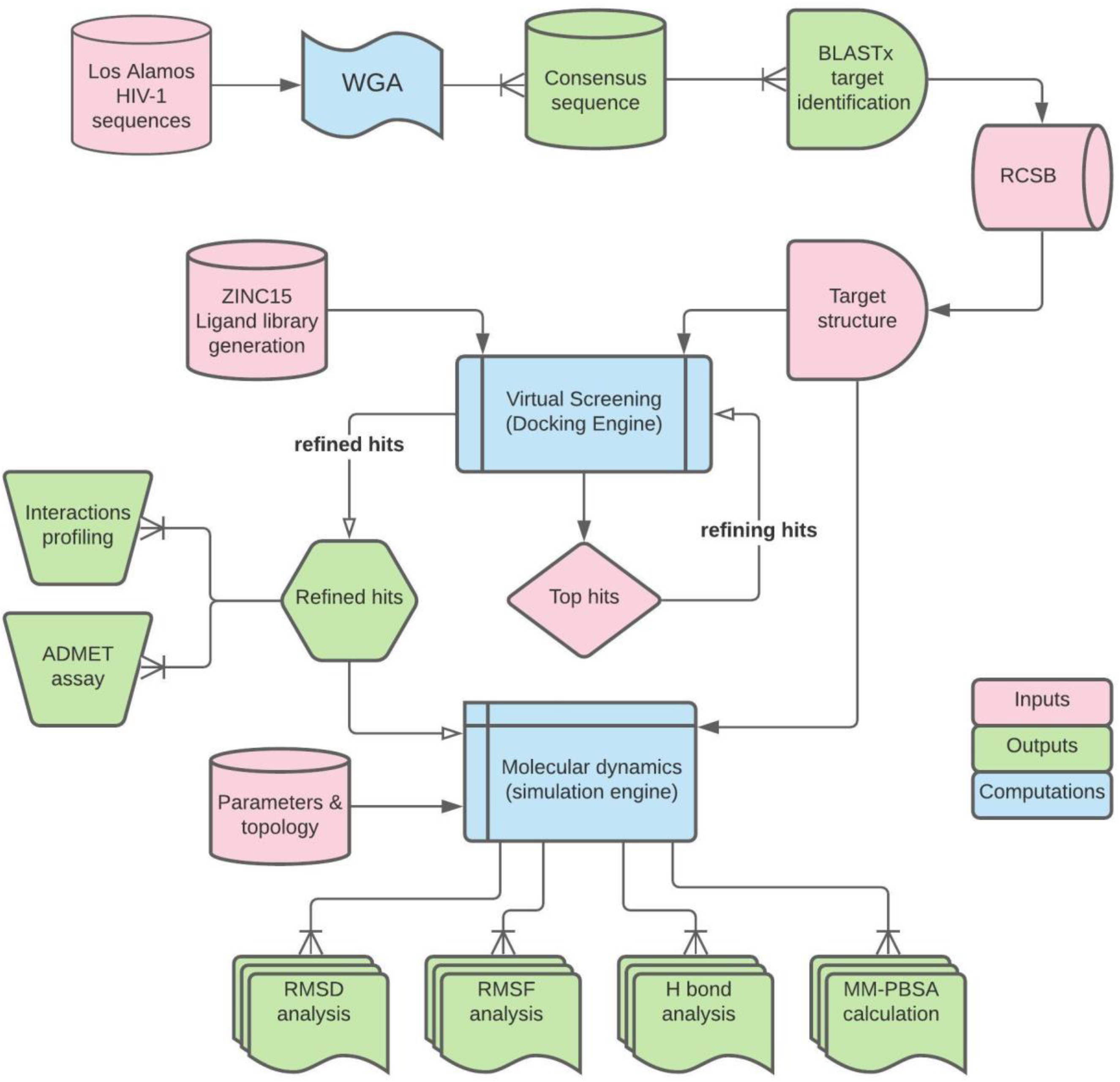
General workflow of the current study.

**Fig 3.**
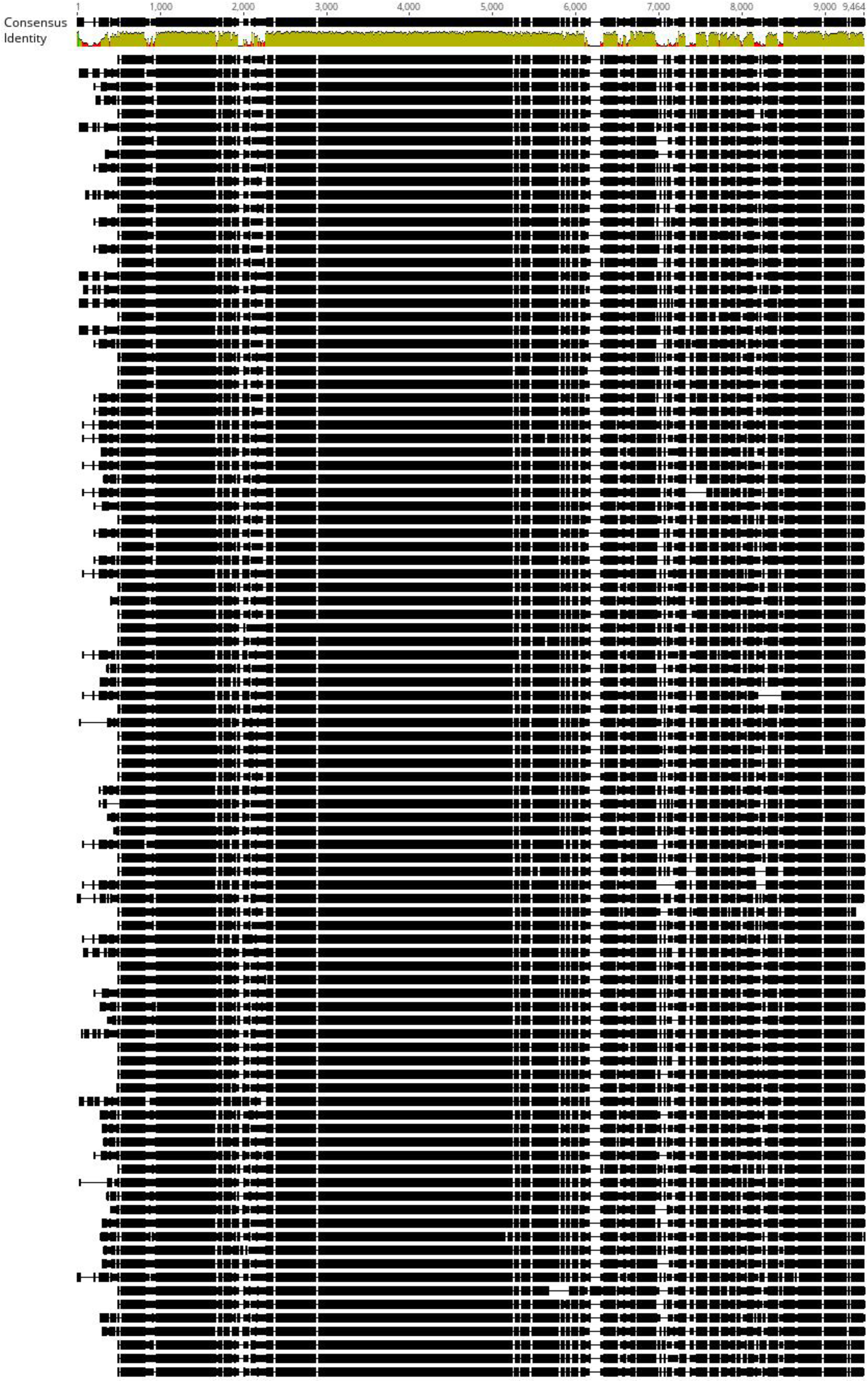
Whole-genome alignment of 98 HIV-1 genomes via progressiveMAUVE algorithm. Identity percentage on top (red for low matches and green for high matches).

**Fig 4.**
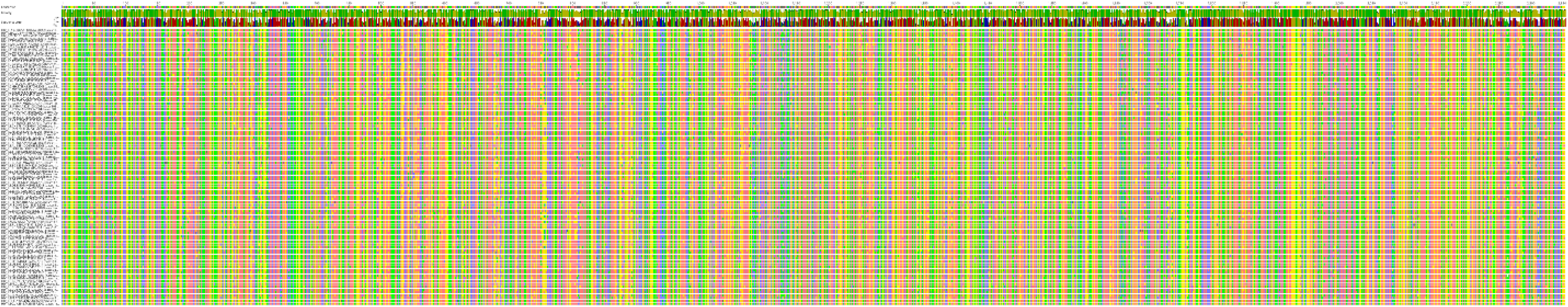
Alignment of the longest highly conserved region within the 98 HIV-1 genomes aligned via progressiveMAUVE (Adenin in pink, guanine in yellow, thymine in green, and cytosine in blue).

The BLASTx results from the consensus sequence have indicated that the highly conserved region belongs to the HIV-1 pol gene (Accession No.: QMX87928.1), hence nominating the functional proteins from the HIV-1 pol gene (RT, IN, and late-phase protease) as a promising target for the development of therapeutics, therefore, further therapeutic screening and analysis were performed on one of the main pol gene products, the reverse transcriptase enzyme [57].

### 2.2. Virtual Screening and molecular docking

The virtual screening of the ligand dataset against the HIV-1 RT enzyme nominated 7 compounds with significantly high affinities (≥ −8.5 kcal/mol), among them only 4 compounds successfully achieved the same affinity in 3 subsequent runs, hence only these 4 compounds were selected and further analyzed, the top docking poses’ for these 4 compounds are shown in Fig 5, a summary of each compound along with their chemical structures is given in Table 2.

**Fig 5.**
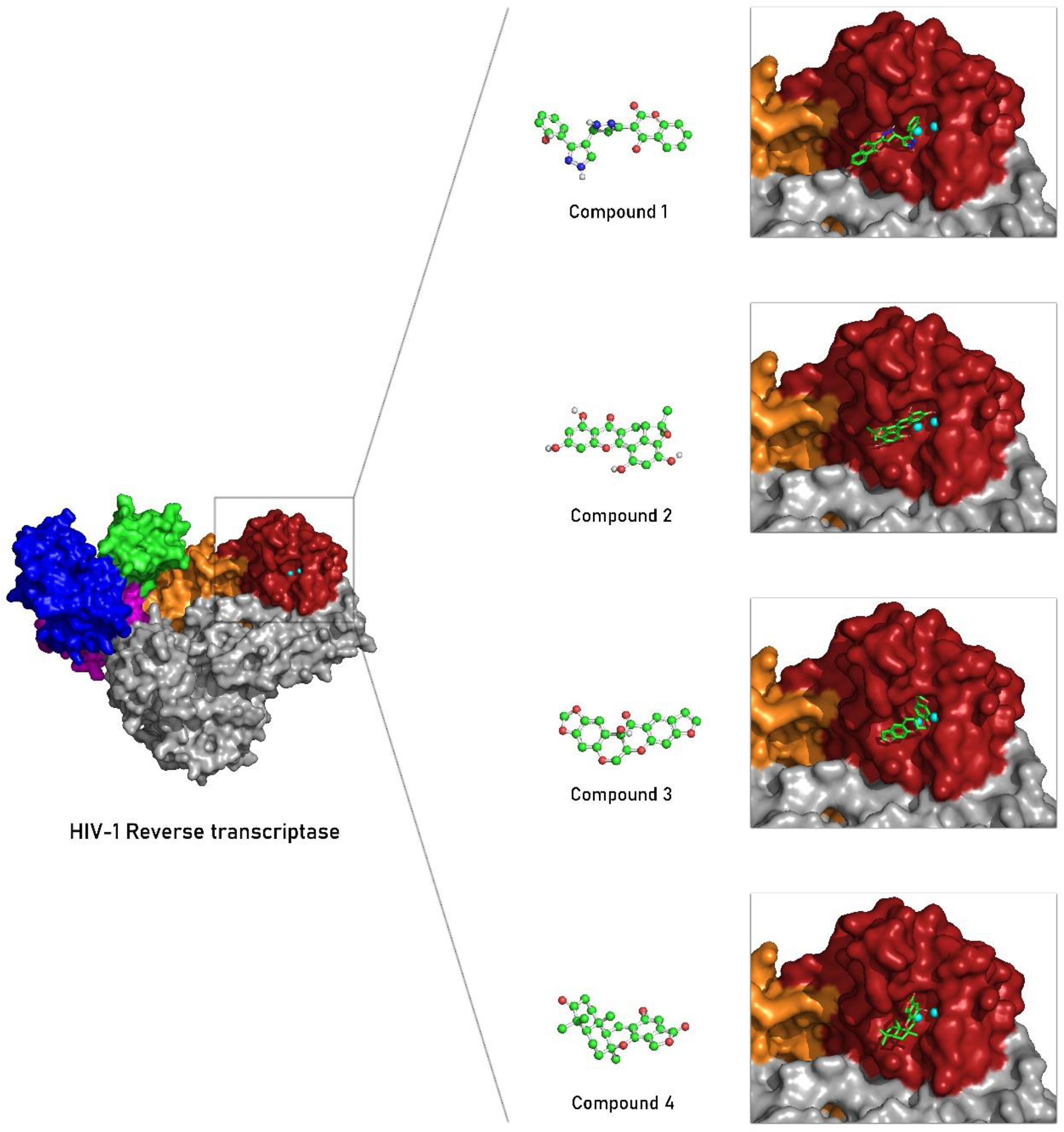
Docking poses’ for the top 4 hit compounds with the highest scores against the HIV-1 RT enzyme. (surface representation on the left). RNase H domain in firebrick red, fingers subdomain in blue, thumb subdomain in green, palm subdomain in magenta, connection subdomain in orange, the cofactor Mn in cyan beads, chain B in gray, and lead compounds with balls and stick representation in light green.

**Table 2.**
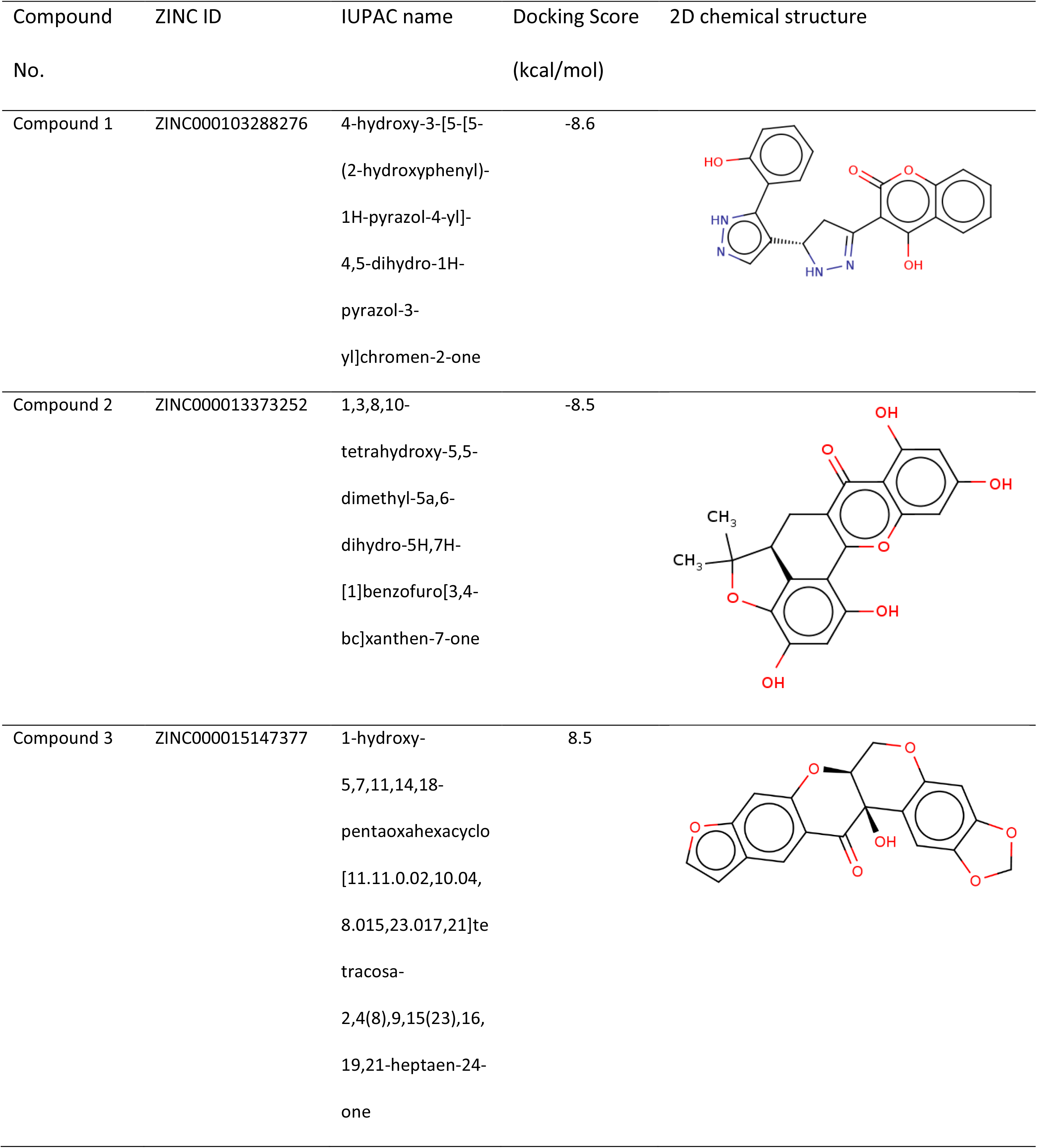
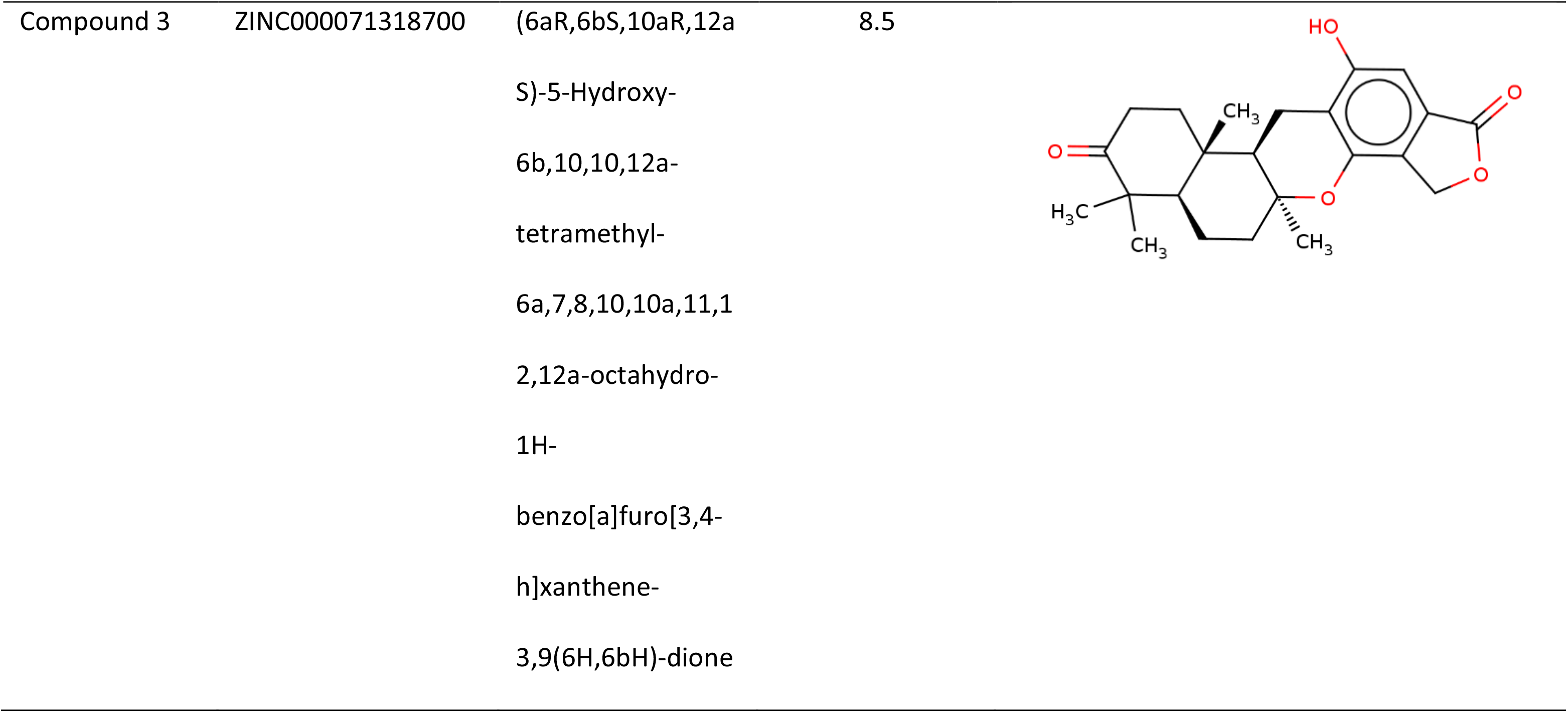
Brief description of the top hit compounds as shown in Fig 5.

### 2.3. Interactions profiling

The interaction between the RT’s RNase H catalytic site with the divalent cation and the top 4 potential lead compounds are (as shown in Fig 5) was closely analyzed and visualized, Fig 6 (a-d) shows all the potential hydrophobic interactions (with yellow dashes) and hydrogen bonds (with magenta dashes) between each residue and lead compound within the RNase H active site, the interacting residues from the RT backbone are further expanded (stick representations in dark wild willow) to visualize the interacting atoms. The right columns in Fig 6 (e-h) shows all the potential interactions between the lead compounds and the cofactor Mn^2+^ cations (cyan beads, dark blue dashes), the residues D443, E478, D498, and D549 (DEDD motif) interacting with the cofactor cations are also expanded to visualize the proximity (sky blue sticks) of the interactions, a summary Table of all the interactions between the lead compounds within the RNase H active site has been listed in Table 3 along with their distances.

**Fig 6.**
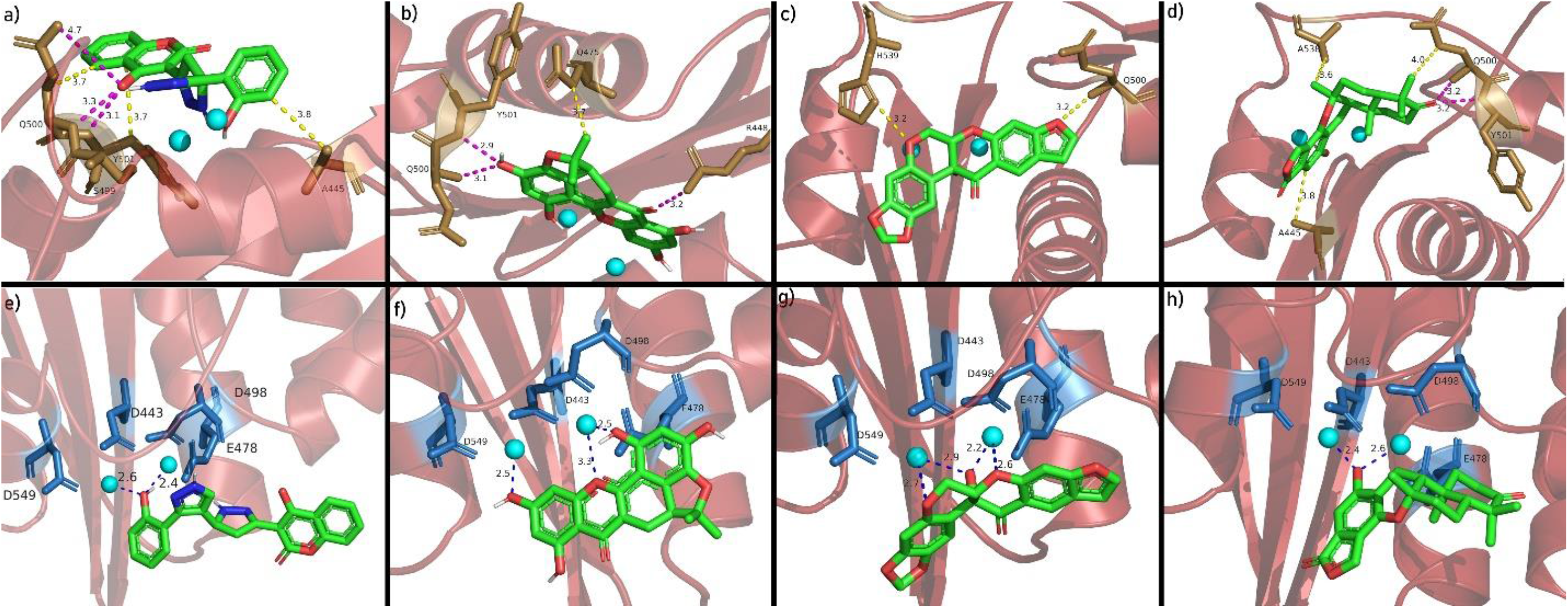
Predicted interactions of each lead compound within the RNase H active site of HIV-1 RT enzyme from their respective top binding pose. All the potential hydrophobic interactions (yellow dashes), hydrogen bonds (magenta dashes), and ionic interactions (dark blue dashes) within the RNase H active site for compound 1 (a & e), compound 2 (b & f), compound 3 (c & g), and compound 4 (d & h) are visualized along with their interaction distances, all lead compounds are shown as green sticks with oxygen atoms colored pink and nitrogen atoms blue. RNase H domain is shown with cartoon representation (firebrick red, semi-transparent), all residues interacting with their respective lead compound are represented with sticks emerging from the RT backbone in dark wild willow color (a-d), the cofactor Mn^2+^ is shown as cyan beads, and the catalytic site residues holding the Mn^2+^ cations (the DEDD motif) is shown in sky blue sticks emerging from the RT backbone (e-h), all measurements are in Å unit.

**Table 3.**
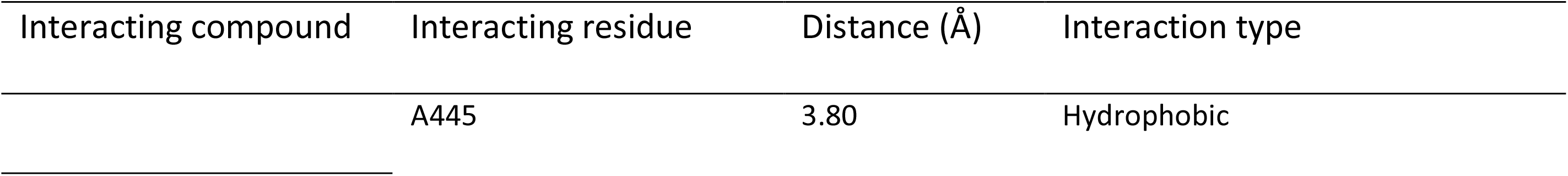
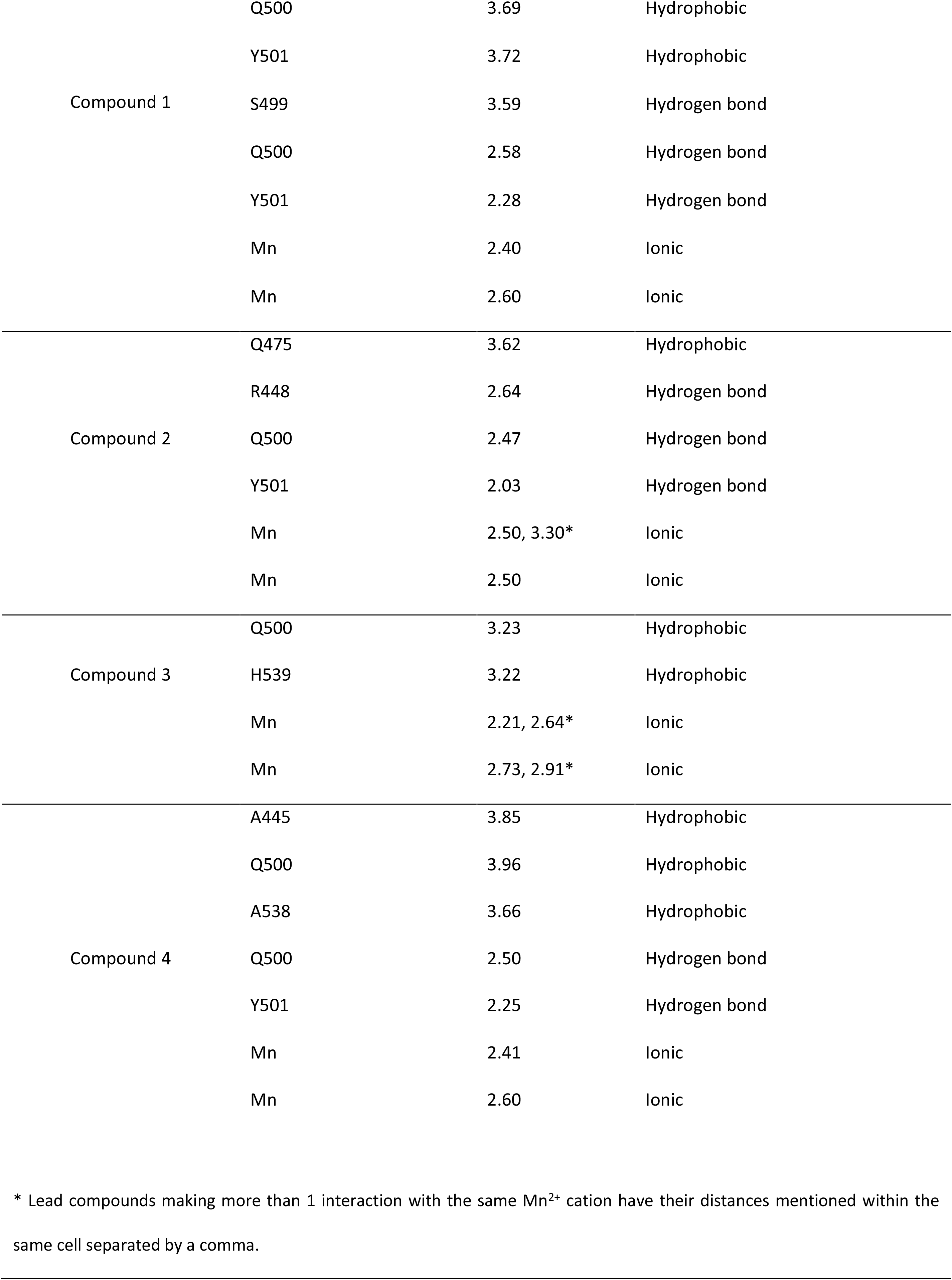
Summary of all the interactions between the RNase enzyme and the respective lead compounds as visualized in Fig 6.

### 2.4. Molecular dynamics analysis

The RMSD for the Cα of the RT from each frame throughout the 50 ns equilibration simulation was extracted and calculated using the 1^st^ frame in each trajectory as the reference point, Fig 7 (a) shows the RT equilibration state throughout the equilibration simulation (plateau in the 40-50 ns interval). Fig 7 (b) shows the Cα RMSF calculated using the same 1^st^ frame as the reference point for each equilibration simulation. An average RMSF plot for the 4 simulations was also calculated by averaging the RMSF of each residue over all of the simulations (Fig 7. c) to analyze the overall RMSF of the RT through the course of equilibration under the simulation conditions.

**Fig 7.**
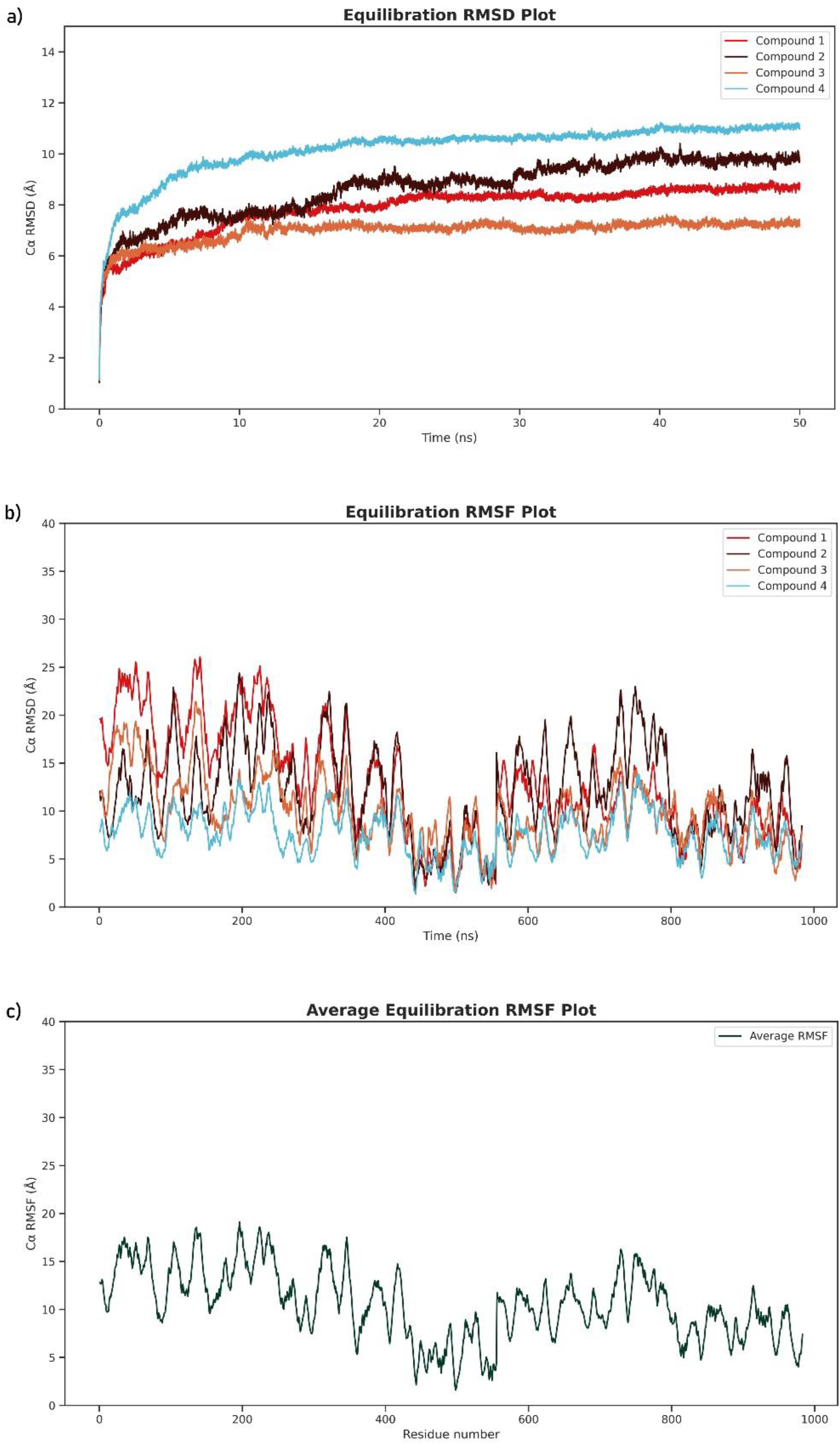
The behavior of the HIV-1 RT enzyme’s backbone through the equilibration simulation. The plot of RT Cα RMSD against time throughout the 50 ns equilibration simulation for each of the RT-lead compound system (a), the plot of RT Cα RMSF against time throughout the 50 ns equilibration simulation for each of the RT-lead compound system (b), and the plot of the average RT Cα RMSF from the 4 RT-lead compound equilibration simulation (c).

The plots for Cα RMSD against time for each RT-lead compound simulation (Fig 8), Cα RMSF for each residue for each RT-lead compound (Fig 9), and the hydrogens bonds formed as a function of time in each RT-lead compound system (Fig 10) throughout the 30 ns production simulation were plotted to subsequently analyze the motion of the lead compound within the simulation system as well as the stability of the RT backbone in presence of the lead compounds.

**Fig 8.**
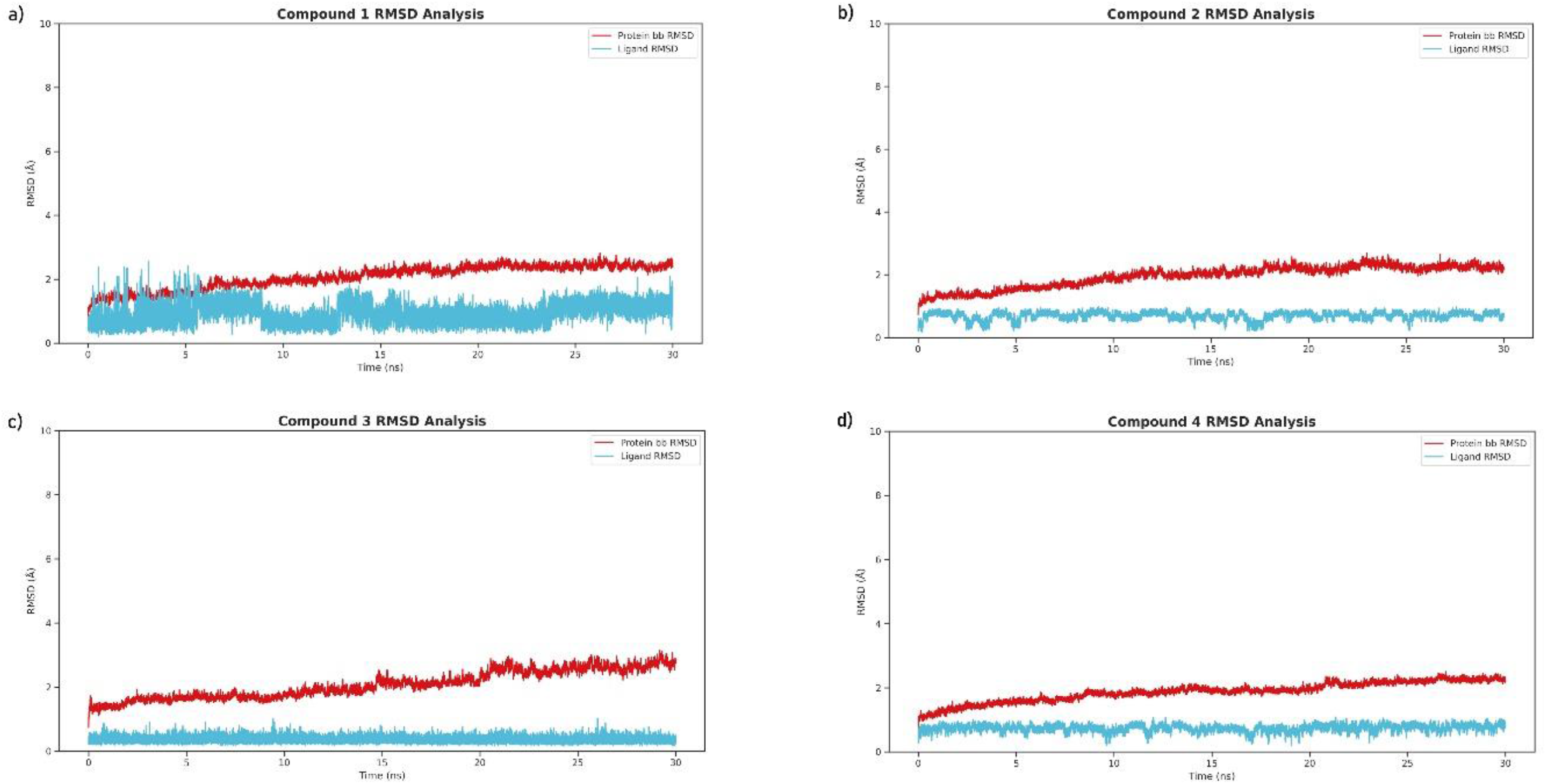
RT backbone RMSD plot with reference to the 1^st^ frame throughout the 30 ns production simulation for compound 1 (a), compound 2 (b), compound 3 (c), and compound 4 (d). The cyan line represents the motion of the lead compound in each system (less variation along the Y-axis implies less deviation from its initial docked position), similarly, the RT backbone RMSD (red line) shows the movement of the RT Cα during the simulation (since it plateaued in Fig 7, most of the motion is random loop movements and/or vibrations).

**Fig 9.**
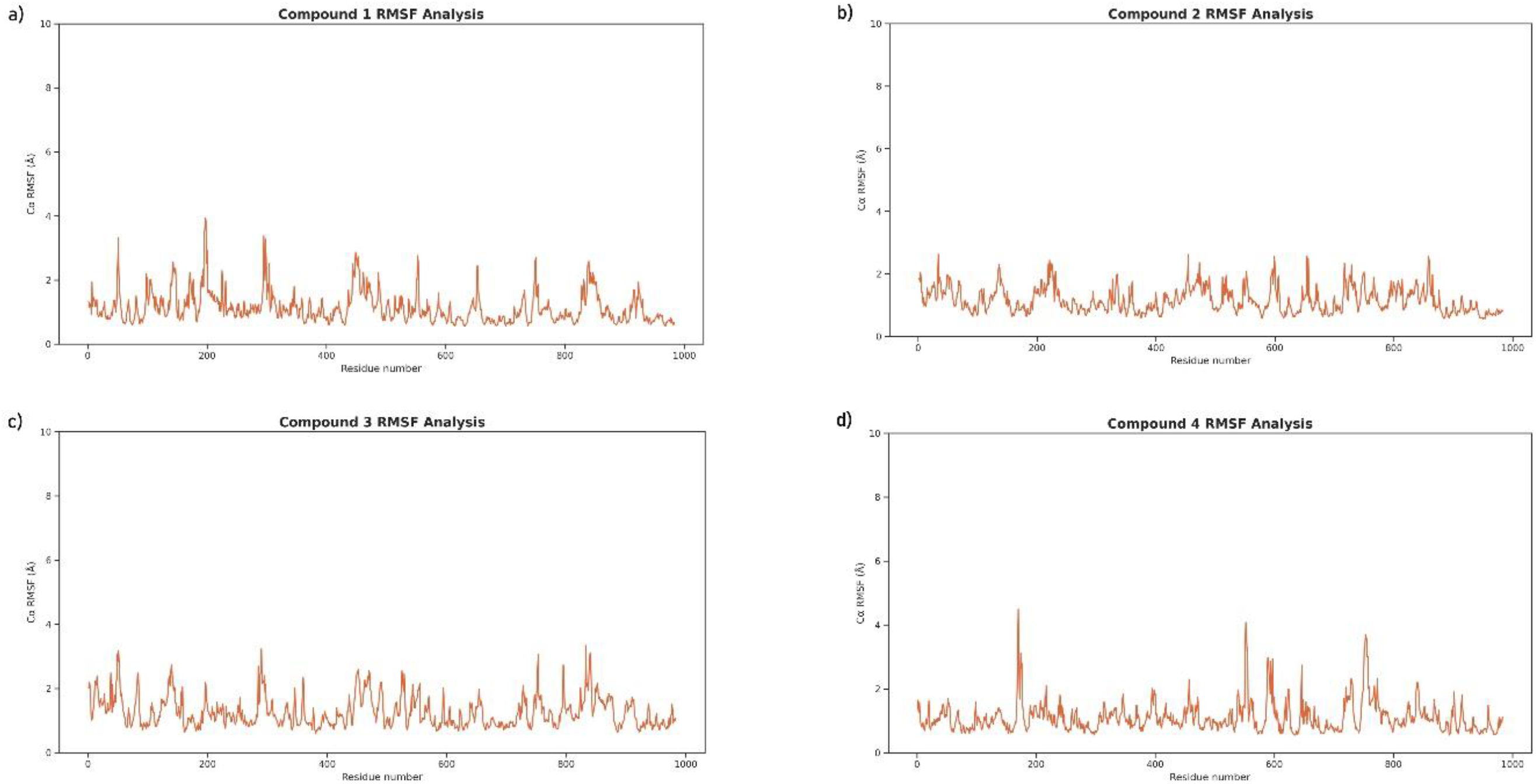
RMSF plot for each residue of RT backbone with reference to the 1^st^ frame throughout the 30 ns production simulation for compound 1 (a), compound 2 (b), compound 3 (c), and compound 4 (d). The peaks (orange line) represent the distance that the corresponding residue (in the X-axis) within the RT moved from its initial state throughout the simulation (lower fluctuation implies a more stable structure).

**Fig 10.**
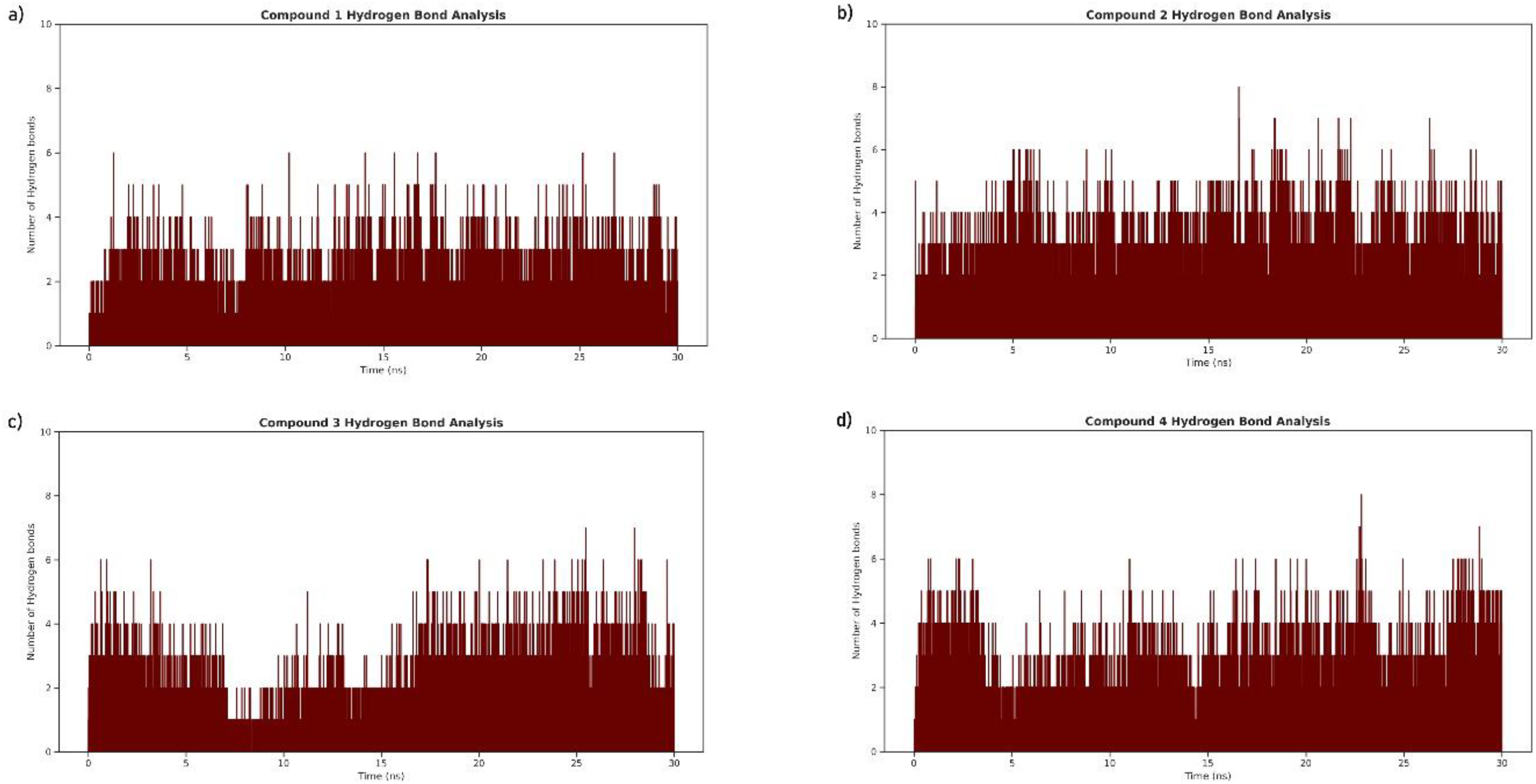
The number of hydrogen bonds formed by each RT-lead compound pair throughout the production simulation for compound 1 (a), compound 2 (b), compound 3 (c), and compound 4 (d). The parameters used to calculate the H bonds were a donor-acceptor distance of less than 3 Å and an angle cutoff of 20° (the minimal threshold for the formation of Hydrogen bonds).

### 2.5. Binding free energy calculation via MM/PBSA

Using the single trajectory approach for MM/PBSA calculation, the 8 energy terms (Δ*E_elec_*, Δ*E_νdw_*, Δ*G_PB_*, Δ*G_SA_*, Δ*G_gas_*, Δ*G_sol_*, Δ*G_pol_*, Δ*G_npol_*) were calculated for the RT enzyme, lead compound, and RT-lead complex separately from each production simulation, and their total sum was used in equation 4 to calculate the binding free energy Δ*G_bind/mmpbsa_*, the sums of each energy term is provided in Table 4 along with their standard deviations, the values of Δ*G_mmpbsa_* indicates the spontaneity of the interaction between the RT and lead compounds (i.e. more negative values implies a more spontaneous reaction). Detailed value for each energy term for the protein, ligand, and complex separately is provided in S5.

**Table 4.** MM/PBSA energy terms for Δ*G_complex_* calculated for each RT-lead pair, energies were calculated from the last 10 ns of the production trajectory using the single trajectory approach.

| $\Delta E_{elec}$ | $\Delta E_{vdw}$ | $\Delta G_{PB}$ | $\Delta G_{SA}$ | $\Delta G_{Gas}$ | $\Delta G_{sol}$ | $\Delta G_{pol}$ | $\Delta G_{npol}$ | $\Delta G_{mmPBSA}$ |
| --- | --- | --- | --- | --- | --- | --- | --- | --- |
| Compound 1 |  |  |  |  |  |  |  |  |
| -28.73±3.76 | -49.67±3.40 | 41.65±2.85 | -5.10±0.12 | -78.40±5.07 | 36.56±2.81 | 12.92±4.48 | -54.77±3.44 | **-41.84±3.90** |
| Compound 2 |  |  |  |  |  |  |  |  |
| -3.68±3.15 | -50.89±2.67 | 30.76±3.84 | -5.38±0.10 | -54.58±3.97 | 25.38±3.83 | 27.08±4.11 | -56.28±2.64 | **-29.20±4.80** |

| Compound 3 |  |  |  |  |  |  |  |  |
| --- | --- | --- | --- | --- | --- | --- | --- | --- |
| -2.27±1.23 | -48.92±2.66 | 13.21±1.20 | -4.90±0.11 | -51.20±3.00 | 8.32±1.18 | 10.94±1.52 | -53.818±2.70 | **-42.88±3.21** |
| Compound 4 |  |  |  |  |  |  |  |  |
| -2.83±1.95 | -54.86±2.56 | 27.58±3.64 | -5.48±0.09 | -57.69±3.02 | 22.11±3.62 | 24.76±4.22 | -60.33±2.54 | **-35.58±4.84** |

± indicates the standard deviations.
All values given energy values are for difference between the complex and sum of protein and ligand, $\Delta X_y = \Delta X_{y(\text{complex})} - (\Delta X_{y(\text{protein})} + \Delta X_{y(\text{ligand})})$ .
All values are in kcal/mol unit.

### 2.6. In silico ADMET assay

The physiochemical properties of the 4 lead compounds are listed in Table 5 along with their drug-likeness results, all 4 lead compounds passed the Lipinksi rule of 5, Ghose filters, Veber filter, Egan filter, and Muegge filter without any violations. The ADMET profiles of each lead compound are also summarized in Table 6 based on the results from admetSAR and ADMETlab.

**Table 5.** Physiochemical and drug-likeness properties of the lead compounds based on the SwissADME results.

| Physiochemical properties | Compound 1 | Compound 2 | Compound 3 | Compound 4 |
| --- | --- | --- | --- | --- |
| Molecular weight (g/mol) | 388.38 | 368.34 | 352.29 | 384.47 |
| No. heavy atoms | 29 | 27 | 26 | 28 |
| No. aromatic heavy atoms | 21 | 16 | 15 | 6 |
| No. rotatable bonds | 3 | 0 | 0 | 0 |
| No. H-bond acceptors | 6 | 7 | 7 | 5 |
| No. H-bond donors | 4 | 4 | 1 | 1 |
| Log S (ESOL) | -4.15 | -4.31 | -3.92 | -4.68 |
| Solubility (mg/mL) | 2.72e-02 | 1.82e-02 | 4.24e-02 | 8.12e-03 |
| Solubility class* | Moderately soluble | Moderately soluble | Soluble | Moderately soluble |
| Lipophilicity (Log P<sub>o/w</sub>)<sup>×</sup> | 2.30 | 2.42 | 2.15 | 3.67 |
| Lipinski rule of 5<sup>†</sup> | Pass (0) | Pass (0) | Pass (0) | Pass (0) |
| Ghose filters<sup>†</sup> | Pass (0) | Pass (0) | Pass (0) | Pass (0) |
| Veber filters<sup>†</sup> | Pass (0) | Pass (0) | Pass (0) | Pass (0) |
| Egan filters<sup>†</sup> | Pass (0) | Pass (0) | Pass (0) | Pass (0) |
| Muegge filters | Pass | Pass | Pass | Pass |

\*Based on the Log S (ESOL) scale, insoluble < -10 < poor < -6 < moderate < -4 < soluble < -2 < very < 0 < highly soluble.
<sup>×</sup> The values are the average of iLOGP, XLOGP3, WLOGP, MLOGP, and SILICOS-IT.
<sup>†</sup> Based on the BOILED-Egg model [58].
<sup>†</sup> Numbers within parentheses indicate the number of violations of the respective filter/rule.

**Table 6.** ADMET profiles of the lead compounds as per results from admetSAR and ADMETlab.

| ADMET properties | Compound 1 | Compound 2 | Compound 3 | Compound 4 |
| --- | --- | --- | --- | --- |
| Absorption |  |  |  |  |
| Gastrointestinal absorption<sup>†</sup> | High | High | High | High |
| Blood-brain barrier permeation<sup>†</sup> | None | None | None | Yes |
| Distribution |  |  |  |  |
| Plasma binding protein<sup>×</sup> | 96.58% | 98.23% | 95.53% | 94.06% |
| Fraction unbound in plasma | 2.52% | 3.30% | 5.60% | 7.13% |

| Metabolism |  |  |  |  |
| --- | --- | --- | --- | --- |
| CYP450 2C9 Substrate | Non-substrate | Non-substrate | Non-substrate | Non-substrate |
| CYP450 2D6 Substrate | Non-substrate | Non-substrate | Non-substrate | Non-substrate |
| CYP450 3A4 Substrate | Non-substrate | Substrate | Non-substrate | Substrate |
| CYP450 1A2 Inhibitor | Inhibitor | Inhibitor | Non-inhibitor | Non-inhibitor |
| CYP450 2C9 Inhibitor | Inhibitor | Inhibitor | Non-inhibitor | Non-inhibitor |
| CYP450 2D6 Inhibitor | Non-inhibitor | Non-inhibitor | Non-inhibitor | Non-inhibitor |
| CYP450 2C19 Inhibitor | Inhibitor | Non-inhibitor | Non-inhibitor | Non-inhibitor |
| CYP450 3A4 Inhibitor | Inhibitor | Non-inhibitor | Non-inhibitor | Non-inhibitor |
| CYP Inhibitory Promiscuity | High | High | Low | Low |

| Excretion<sup>x</sup> |  |  |  |  |
| --- | --- | --- | --- | --- |
| T<sub>1/2</sub> (hours)<sup>†</sup> | < 3 (0.349) | >3 (0.52) | < 3 (0.113) | >3 (0.70) |

| Toxicity |  |  |  |  |
| --- | --- | --- | --- | --- |
| Rat acute (LD50, mol/kg) | 2.43 | 2.38 | 2.41 | 2.71 |
| TP* (pIGC50, ug/L) | 0.46 | 1.04 | 0.57 | 1.38 |
| Acute oral (LD50 mg/kg) <sup>‡</sup> | III | III | III | III |
| Carcinogenicity | None | None | None | None |

<sup>x</sup> Based on predictions of ADMETLab 2.0 [59].
<sup>†</sup> Probability of half-life being greater than 3 hours is given within parentheses, below 0.5 was considered to have T<sub>1/2</sub> < 3.
\* *Tetrahymena Pyriformis* toxicity.
<sup>‡</sup> Class I ≤ 50 mg/kg, class II > 50 mg/kg, class III > 500 mg/kg, and class IV > 5000 mg/kg.
The SMILES notation of the lead compounds was used as the input to calculate each of the properties listed in the table is provided in S6.

## 3. Discussion

Ever since its emergence, HIV infections have kept on harvesting lives, while modern anti-viral and HAART therapies provide some relief and support for the patients, it comes with its disadvantages and limitations. As mentioned in the introduction, this study aimed to perform extensive in silico analysis to discover and evaluate potent novel inhibitors of HIV-1 replication within the host by targeting the most coherent target, providing therapeutic options for the patients while at the time accelerating the drug development processes by providing potential leads.

The mutation rate for HIV-1 is reported to be 10^−4^ to 10^−2^ mutants/clones and with the estimated production of 10^9^ virions/day within an infected individual, the virus mutates quite efficiently to develop resistance and/or evade the immune system [60]. However, not all of these mutants are expected to survive and replicate as mutations to some genes could be lethal, hence the most coherent target for drug discovery and development efforts would be the phenotypes that mutate less frequently as their chance of developing resistance or evasion are lower than their highly mutating counterparts, to determine such regions within the HIV-1 genome, the comparative genomics approach was used, comparative genomics considers the correlations and differences between the genotype (genome) of closely related species or even different variants of the same specie to answer the reasons behind their characteristic phenotypes, this method has been widely used to determine resistance genes in several bacterias in the past [61–63]. In this study, we utilized 98 high-quality HIV-1 sequences from the Los Alamos database and performed whole-genome alignment with MAUVE to determine the regions within the HIV-1 that genome that shows the highest level of consensus among all of the 98 sequences, as visualized in Fig 3, a genomic fragment of around 2.4 kb long was the longest genomic fragment (the green bars on top of the sequences in Fig 3) with the least variation among all of the 98 sequences, BLASTx was utilized to determine what this genomic region encoded for and the result from BLASTx deducted the pol gene, which encodes for 3 important functional proteins with the HIV-1, the viral reverse transcriptase, integrase, and late-phase protease, all of which are coherent targets for drug targeting.

The RT enzyme being among the most extensively studied proteins of HIV-1, this study aimed to discover and analyze potent targets inhibiting the RT enzyme itself, given that several NNRTI and NRTI has already been approved by the FDA for use and since all of them function by inhibiting the DNA polymerase activity of the RT, the aim of the study focused specifically on discovering and analyzing potent RT RNase H inhibitors, which is equally critical for the viral replication as its polymerase activity [64–67].

Virtual screening has been used extensively in recent years to discover novel therapeutics and/or repurpose existing drugs to new targets, in summary, it can be viewed as a form of batch molecular docking where an attempt is made to position each compound within a library of compounds to the best position it can have within a specified region of a protein, usually an active site or a receptor’s binding domain, such that it would attain maximum affinity towards it, this method is more efficient, faster, and cheaper than the high throughput screening (HTS) approaches [55,68,69]. In this study, a library of 94,545 small molecules stable at physiological pH with a charge of 0, −1, and −2 (since the HIV-1 RNase H active site contains positive Mn/Mg cofactors) was generated with their weights limited to be in the range of 350-500 g/mol (based on the size of the RT’s RNase H active site pocket). The virtual screening result initially reported 7 hits with affinity scores below −8.5 kcal/mol, upon 3 subsequent repetitions, only 4 of these compounds successfully reproduced the same affinity scores in the same pose within the RNase H active site (referred to as Compound 1, 2, 3, and 4 in this study, also briefly described in Table 2) and hence were considered for further analysis. The scoring function of AutoDock Vina is based on a 190-complex training set only, combining this with the fact that docking experiments don’t consider the water, salts, temperature, pressure, and other cellular factors, the binding affinity from docking experiments are not absolute, hence, molecular dynamic and MM-PBSA binding free energy approaches were integrated to both validate the docking pose and estimate the binding free energies of the respective lead compounds. Upon closely investigating the binding pose of each of the lead compounds within the RT’s RNase H active site (as shown in Fig 6), the binding poses are reasonable as all the lead compounds form several hydrogen bonds except for compound 3 only, all the leads also happen to be in less than 4 Å proximity to form hydrophobic and ionic interactions with the active site residues and the active site Mn cofactor which provides some basis to the potential these lead compounds have in being actual potent inhibitors [70,71].

Molecular dynamics provide a great opportunity to explore and analyze the behavior of biomolecules at nano-scale levels, which makes it an indispensable tool for drug discovery, as the interactions a drug could make under simulation conditions designed to replicate the physiological cellular conditions are more likely to represents its interactions in *in vitro* and *in vivo*, and therefore dictate its activity and side effects [72,73]. In both X-ray and Cryo-EM (cryogenic electron microscopy), the protein structures are not determined in physiological conditions but rather in extremely low temperatures (often liquid Nitrogen temperature), hence the first step in simulating near-actual physiological conditions is to equilibrate the protein into its native state under the physiological temperature [74]. Performing equilibration of proteins in a single step is often ill-advised as it can cause irregularities and clashes within the structure resulting in unexpected behaviors and biases during downstream analysis, hence, in this study, each of the 4 RT-lead compound systems was minimized and equilibrated in 5 steps, first, the energy of the system was minimized for 2 ns, then the RT-lead complex along with the Mn^2+^ cations was restrained to allow the water and salt ions to equilibrate around the RT-lead complex for 5 ns, then the restraints on the protein’s side chains was released to allow them to equilibrate for 10 ns, following this step was the whole system equilibration for 50 ns with only the Mn^2+^ cations restained to avoid it from drifting away, and finally, 30 ns simulation with no restraint was performed from which the trajectory was collected for analysis [75]. To ensure that the RT enzyme was well-equilibrated before performing the production simulation, the RMSD of the RT’s Cα was plotted throughout the 50 ns equilibration simulation to ensure it reached a plateau, which happened as seen in Fig 7 (a), the RT enzyme in all of the 4 systems reached a plateau after the 30 ns time-lapse which meant the production runs would provide accurate and less biased results for the behavior of the lead compounds. The HIV-1 RT enzyme is a large complex with many loop regions, which unlike α-helix and β-sheet conformations have no absolute stable conformation, hence they often move randomly throughout the simulation resulting in higher overall RMSD and RMSF, this phenomena can be noticed in Fig 7 (b and c), where residues close to the C- and N-terminal, i.e. the fingers subdomain (residue 1-85 & 118-155, consisting of several loops) and the palm subdomain (residue 86-117 & 156-236, containing some long loops) has higher RMSF values in each of the 4 simulation systems. The RMSD plot of compound 1 (Fig 8a) is definitively a negative result as compound 1’s backbone (cyan line) appear to deviate ≈ 2 Å throughout the simulation, this implies that on average, compound 1 had moved ≈ 2 Å from its initial docked position, which can be interpreted into it leaving the active site, thereby possessing no potent affinity towards the RT enzyme’s RNase H domain, compound 2 and 4 (Fig 8 b & d, cyan line) however, deviated less 1 Å which are ideal values to say that they stayed mainly within the active site and therefore possess an affinity to the RNase H active site, compound 3 (Fig 8c, cyan line) on the other hand appears to possess the best affinity as it barely moved 0.5 Å from its initial docked pose. The red lines in Fig 8 represent the RT’s Cα RMSD throughout the production simulation, ensuring that the RT enzyme itself didn’t undergo any unusual conformational change (fragment, aggregate, change in conformation, etc due to the presence of the ligand). The RMSF plots in Fig 9 were also plotted to ensure that the RT enzyme structure was not compromised in any of the simulation systems, since the enzyme structure was already equilibrated for 50 ns prior (their RMSD graphs reached a plateau in Fig 7), no high peaks are to be expected like those of Fig 7 (above 10 Å RMSD), Cα RMSF below 5 Å is mainly contributed by the vibration effect and random movement of the loop regions [76]. Fig 10 shows all the potential hydrogens between each lead compound and the residues within 3 Å range from them at a maximum cut-off angle of 20°, each lead compound retains at least 3 hydrogen bonds on average throughout simulation which is relatively similar to the hydrogen bonds predicted using PyMol (Fig 6 and Table 3) except for compound 3, which was predicted to form no hydrogen bonds, however, even in Fig 10 (c) compound 3 doesn’t form stable hydrogen bonds until it passes the 18 ns mark, considering its RMSD of around 1 Å (Fig 8c), it probably rotated around itself to some degree and/or got closer to other nearby residues to form these hydrogen bonds and even then, it made on average fewer hydrogen bonds than the other lead compounds.

To determines the spontaneity or propensity of a drug candidate at a given (physiological) temperature, its binding free energy toward its biomolecular target is used as an objective quantity in the selection process. Binding free energy calculations are quite challenging both *in vitro* and *in silico*, however, *in silico* routes are less labor-intensive and more computationally expensive, in this study the fast and accurate MM-PBSA protocol was used on the last 10 ns (where the RMSD had reached a plateau and were stable) of each simulation system to estimate their binding free energies, a total of 8 energy terms was calculated for each of the RT enzyme, the lead compound, and the RT enzyme in complex with the lead compound. Equation 4 was used to calculate Δ*G_mmpbsa_* which was also approximated to be equal to the respective Δ*G_bind_* as mentioned in equation 5 (and justified by the former reasons) [77,78]. The binding energies for each of the compounds (in Table 4) are promising with the best result from compound 3 (which also had the lowest overall RMSD) followed by compound 1, compound 4, and finally, compound 2, all the estimated Δ*G_bind_* values being negative can conclude that each of these compounds can spontaneously bind to the RT’s RNase site, however, given the RMSD plot of compound 1 (Fig 8a), it is unlikely that this value is true for the RNase H active site since it drifted from the active site ≈ 2 Å, hence, its binding free energy is to some region ≈2 Å away from the RNase H active, therefore, from the molecular dynamics simulations and MM-PBSA calculations, we can conclude that compound 2, 3, and 4 has a significant affinity to the HIV-1 RT’s RNase H active site and thereby could potentially hold some inhibitory activity.

Evaluation of the physicochemical, pharmacological, and ADMET profiles of the lead compounds has been used extensively in the very early stages of drug discovery to accelerate the conversion of leads into qualified development candidates, most of these evaluations are performed *in vitro* which costs a lot of time and resources, however, with the developments in the field of machine learning and quantitative structure-activity relationship (QSAR), these experiments could be performed in silico with statistically significant results, this study utilized the pre-trained algorithms of 3 different ADMET assessing webserver to conclude the physiochemical, pharmacological, and ADMET profiles of the lead compounds analyzed [79–81]. Since compound 1 had shown to not poses proper inhibitory potentials from previous analyses, further interpretations focused strictly on compounds 2-4. As given in Table 6, each of compounds 2-4 exercised high gastrointestinal absorption levels which are critical for any drug as they have to be absorbed into the body to exert their effect, among the lead compounds, only compound 4 showed the potential to permeate blood-brain barrier which makes it’s a valuable lead as HIV-1 is known to cross the blood-brain barrier and infect target cells causing neurocognitive disorders and brain damage in the processes [82,83]. As for the distribution of the compounds, the most important parameter is the fraction of the compound unbound in the plasma, ideally, the higher is better as the unbound fraction is the active drug available to exert its function (>5% is a decent threshold), among the lead compounds, both of compound 3 and 4 provide accepTable unbound fractions for a therapeutic agent [84,85].

Cytochrome P450 (CYP450) enzymes are essential for the metabolism of many drugs, around 50 isoforms of CYP450 enzymes are identified in humans, however, 90% of drugs are known to be metabolized by 6 of these isoforms (CYP1A2, CYP2C9, CYP2C19, CYP2D6, CYP3A4, and CYP3A5), the overall performance of the lead compounds was evaluated by CYP inhibitory promiscuity parameter (Table 6), the classification of high and low refers to whether the respective compound does or does not possess substantial CYP450 inhibitory promiscuity, lower CYP inhibitory promiscuity indicates less risk of clinical drug-drug interaction and better/higher metabolism (and therefore excretion as well) of the drugs, among the lead compounds reported in this paper, both compound 3 and 4 exhibits low CYP inhibitory promiscuity making them ideal choices for drug candidacy [86,87]. The excretion of the lead compounds was evaluated based on their half-life (T_1/2_) within the body, this parameter indicates the time it takes for the lead compounds to reach half of their initial concentration in the body once administered. While there is no ideal benchmark for T_1/2_ of therapeutic drugs, 12-48 h are often considered ideal as shorter T_1/2_ would require more frequent intake of the drug to maintain its activity and longer T_1/2_ would cause accumulation of the drug and prolonged side effects [88]. Among the lead compounds reported in this paper, compounds 2 and 4 exhibit T_1/2_ greater than 3 hours (although with a lower probability for compound 2) whereas compound 3 exhibited T_1/2_ lower than 3 hours, further nominating compound 4 as a potent drug candidate. Table 6 also reports the toxicity of the lead compounds based on pre-trained regression models by admetSAR, admetSAR database contains 5 quantitative regression models with high predictive accuracy as their models are trained with over 210,000 ADMET annotated data points for more than 96,000 unique compounds with 45 kinds of ADMET-associated properties curated from literature [89]. According to its toxicity predictions based on rat acute toxicity and *Tetrahymena Pyriformis* toxicity, compound 4 is the best performing lead, ideally, the higher the LD50 and pIGC50 for rat acute toxicity and *Tetrahymena Pyriformis* toxicity respectively, the higher the dose it would require to attain a toxic effect, hence providing a wider range for a therapeutic dose [90]. None of the lead compounds were predicted to exhibit any carcinogenic activity. The acute oral toxicity parameter was predicted based on a model trained on 12,204 diverse compounds with their LD50 classified based on the criterion of US EPA for all drugs, all of the lead compounds reported in this paper were predicted to have an acute oral toxicity between 500-5000 mg/kg, which a implies that doses below this range are unlikely to result in any acute oral toxicity, hence each one of them are suitable for oral administration.

Considering all the in silico analysis reported in this paper and the lead selection criteria applied, compound 4 is the best performing lead compound, with a docking score of −8.5 kcal/mol, several hydrophobic, hydrogen bond, and ionic interactions with active site residues of the HIV-1 RNase H and the cofactor Mn^2+^ cations, it had only shown less than 1 Å deviation from its initial docked pose throughout the 30 ns molecular dynamic simulation, and attained a binding free energy of ≈ −35.58±4.84 kcal/mol with a near-ideal ADMET profiles, a video of the production simulation of the RT with compound 4 is provided in S7. Compound 4, with the IUPAC name (6aR,6bS,10aR,12aS)-5-Hydroxy-6b,10,10,12a-tetramethyl-6a,7,8,10,10a,11,12,12a-octahydro-1H-benzo[a]furo[3,4-h]xanthene-3,9(6H,6bH)-dione is also known in the literature as Phomoarcherin B. Phomoarcherin B (PubChem CID 52952104) is a natural compound found in the endophytic fungus *Phomopsis archeri,* it was first isolated and characterized as a pentacyclic aromatic sesquiterpene via spectroscopic analysis by Hemtasin et al. (2011) and tested for antimalarial and anticancer activities on cholangiocarcinoma cell lines [91]. It was also featured in 2 different reviews for the same characteristics, however, no in vitro or in vivo assay has been performed regarding its antiviral or RNase H inhibitory activity, making it a potential novel lead that could inhibit HIV-1 replication by inhibiting its RNase H activity [92,93]. Compound 2 and 3 also exhibit potential lead-like characteristics with only some bad scores on their ADMET profiles, hence further *in vitro* assays are necessary to confirm their potency as potential HIV-1 RNase H inhibitors.

Given the potency of Phomoarcherin B as a strong anti-RNase H agent, additional docking studies was performed against other viral RNase H enzymes, mainly the Feline immunodeficiency virus (FIV) RT RNase H domain, the RT RNase H domain of the Moloney murine leukaemia virus (MLV) and RNase H of Bacteriophage T4. The docking trials yielded promising results with affinities of −9.0, −8.3, and −8.1 respectively. This results further open the doors for further studies to confirm the potency of Phomoarcherin B as a general viral anti-RNase H agent. The top docking pose of Phomoarcherin B with the former enzymes is shown in Fig 11 and more details are provided in S8.

**Fig 11.**
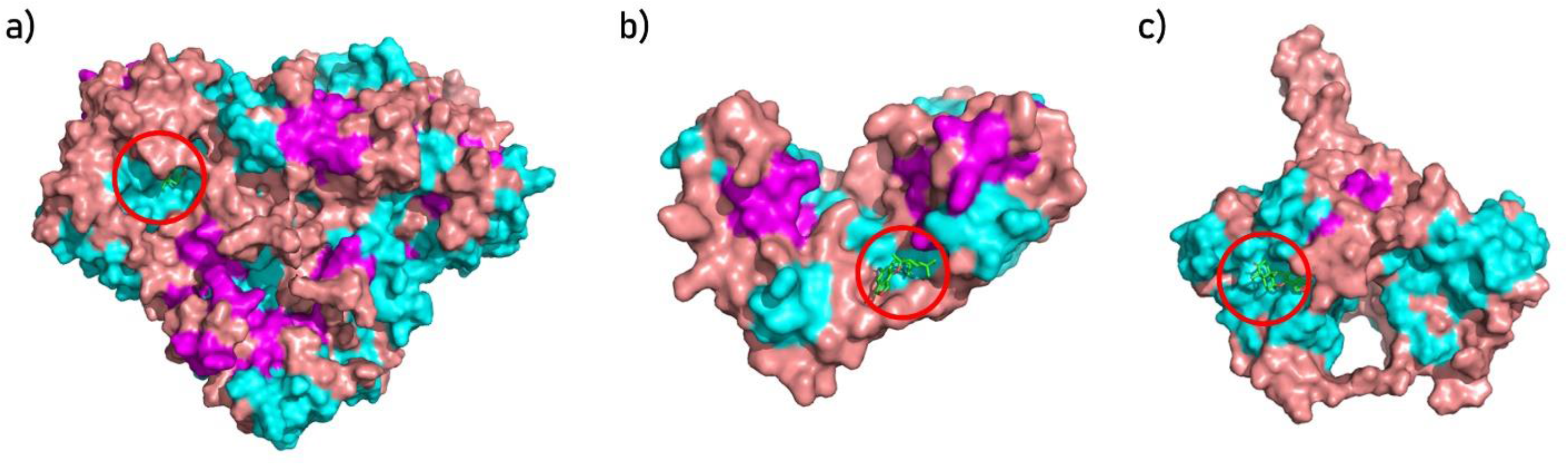
Top docking pose of Phomoarcherin B (red circle) with the RT of FIV (a), RT of MLV (b), and RNase H of the Bacteriophage T4 (c). Phomoarcherin B represented as green sticks and bonds, proteins in surface representation colored based on their (helices in cyan, sheets in light pink, and loops dry violet).

## 4. Materials and Methods

### 4.1. Whole-genome alignment and BLASTx

A total of 98 HIV-1 complete genomic sequences was retrieved from the Los Alamos HIV sequence database [19]. The sequences were manually selected such that only 2 sequences (where applicable) were chosen from each country and the dataset included all the geographic regions available (however, this trend was not strictly followed as some countries like the US received higher coverage for its size and the number of high-quality sequences deposited whereas other countries like India despite its size, due to lack of abundant high-quality sequences deposited, received lower coverage). Whole-genome alignment was performed via progressiveMAUVE algorithm with match seed weight set to automatic calculation, minimum Locally Collinear Blocks set to default (3 times the minimum match size), progressive Muscle (v3.6) was selected for as gap aligner for each LCB, and minimum island size, maximum backbone gap size, minimum backbone size were set to 50 [94,95]. The list of all the sequences used for the alignment is included in supplementary materials 1 (S1) and the whole genome alignment result is included in S2. The alignment result was visualized in Geneious Prime (v2020.1) and the highest conserved continuous region with no gaps in the alignment was excised from the alignment and visualized separately in-depth [96]. A consensus identity sequence from the conserved fragment was generated using Jalview and submitted to NCBI BLASTx with the default parameters (max target sequences 100, expected threshold 0.05, word size of 6, max match in a query range 0, matrix BLOSUM62, gap costs for existence 11, an extension of 1, and compositional adjustments via conditional compositional score matrix adjustment), the alignment for the excised fragments are provided in S3 and the consensus sequence is provided in S4 [97–99].

### 4.2. Molecular docking based virtual screening

The experimentally determined X-ray diffraction structure of HIV-1 RT with PDB ID 3IG1 was retrieved from the Research Collaboratory for Structural Bioinformatics (RCSB) website [30,100]. The missing residues from the structure were added via the PyMol’s builder plugin (open source v2.5.0), the loop regions where the residues were added was refined using MODELLER (v10.1) [101–103]. The structure was then cleaned from all heteroatoms except for the cofactor atoms, polar hydrogens were added where necessary, and the Kollman charge model was applied [104]. A library of 94,545 annotated anodyne small molecules (ligands) stable at physiological pH and having a charge of 0, −1, or −2 was generated from the ZINC15 database [105]. A grid box with a size of 25 Å X 32 Å X 32 Å along the X, Y, Z-axis was calculated (a box around the RNase H active site). Virtual screening was performed with HIV-1 RT structure against the ligand dataset within the grid box calculated at exhaustiveness of 64 (indicates the intensity of the search) via AutoDock Vina (v1.1.2) [106]. The top 7 molecules with the highest affinity scores were screened again with the same configuration but with exhaustiveness of 256, compounds that successfully reproduced their scores in the same pose were retained for further analysis.

### 4.3. Protein-ligand interactions profiling

The best dock pose of the top hit ligands was loaded with the HIV-1 RT to PyMol and all the residues within 4 Å from the lead compounds were visualized (i.e. all potential, hydrophobic interactions, hydrogen bonds, and ionic interactions) and evaluated, the manually predicted interactions were also cross-validated with the TU Dresden’s Protein-Ligand Interaction Profiler (PLIP) webserver and only overlapping interactions were considered and visualized.

### 4.4. Molecular dynamics simulation

The molecular dynamics simulation was performed with the University of Illinois’s Nanoscale Molecular Dynamics (NAMD v2.14 CUDA) tool [107]. OPLS-AA/M force field from William L. Jorgensen research group was utilized to generate the topology and parameter files for the RT enzyme, the same force field was used to parameterize the ligand molecules as well (via LigParGen server with 1.14 CM1A charge model) [108–111]. Each pair of RT-ligand complex was immersed in a square box with explicit TIP3P water such that such a distance of 5 Å was maintained between the edge of the box to the RT-ligand complex along each axis, the system was neutralized with Na^+^ and Cl^-^ ions and their final concentration was maintained at 0.15 mol/L (physiological salt concentration). The system was minimized for 2 ns to reach its lowest energy relaxed state from the X-ray diffraction state, the system was then equilibrated for 5 ns at 310 K with periodic boundary conditions, Langevin dynamics, Particle Mesh Ewald (PME) for electrostatics, and Langevin piston (at 1 atm pressure) with the RT-ligand complex constrained to allow the water and ions equilibrate around the complex, this step was followed by 10 ns equilibration with the constraints on the protein side chains released to allow the side chains to relax. The system was then subjected to 50 ns equilibration with constraints only on the cofactor Mn^2+^ cations to allow the system to reach its equilibrium while maintaining the cofactor in the active site. The root mean square deviation (RMSD) and root mean square fluctuation (RMSF) of RT’s Cα from equilibration simulation was calculated using the 1^st^ frame as the reference point to monitor the RT’s behavior under the simulation system. Finally, a 30 ns production simulation with no constraints was performed from which the trajectory was collected for analysis.

Throughout the simulation, the outputs were written to the trajectory every 1 ps, and RMSD of RT’s Cα and the lead ligand as a function of time elapsed was plotted, RMSF of the Cα for each residue of the RT throughout the production simulation was also plotted. The number of hydrogen bonds between the RT enzyme and the respective lead ligand throughout the production simulation was also plotted (with the donor-acceptor thresholds set to < 3 Å and angle cutoff set to 20°), all the statistical analysis and visualizations was performed using the *Matplotlib* and *Seaborn* libraries [112,113].

### 4.5. Binding free energy calculation via MM/PBSA

The binding free energy (ΔG_bind_, Gibbs free energy) between the lead compounds and RT enzyme was calculated from the last 10 ns (stable RMSD interval) for each simulation system (compromising of 1001 snapshots) via the MM/PBSA single trajectory protocol, the formula in equation 2 was followed to calculate the energy terms for each of the RT enzymes, lead compounds and RT-lead complex using the CaFE plugin (v1.0) in VMD. Finally, equation 4 was used to calculate the ΔG_bind_ [114,115].

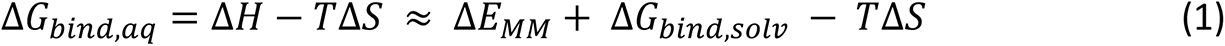

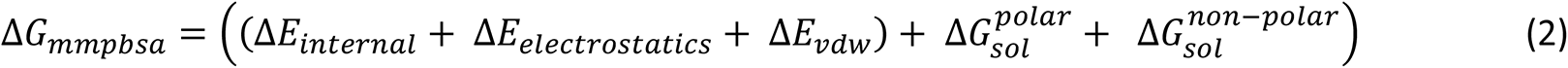

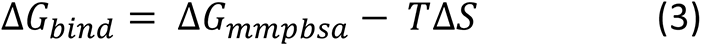

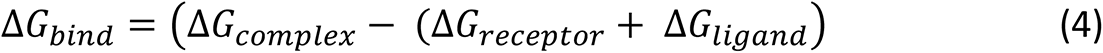

Where Δ*E_internal_* = (Δ*E_bond_* + Δ*E_angle_* + Δ*E_torsion_*), the sum of (Δ*E_internal_* + Δ*E_electrostatics_* + Δ*E_νdw_*) aka Δ*E_MM_* represents the changes of the gas phase molecular mechanic energy, and −*T*Δ*S* is the product of temperature and the change in the conformational entropy upon binding. One of the limitations of in silico end-point binding free energy calculation methods is the difficulty in capturing the changes in configurational entropy involved with the association of the protein with the ligand, this step requires adequate sampling of all the relevant movements of the ligand to the corresponding protein, however, experimentally it demands extensive computation and often provides highly inaccurate results, hence, most in silico approximate binding free energy estimator protocols tend to drop the conformational entropy terms in Δ*G_bind_* calculations, which was also followed in this study, therefore, the Δ*G_bind_* values were calculated using equation 5 to calculate the approximate Δ*G_bind_*, to compensate for this, the gas phase (Δ*G_gas_*) energy calculation was performed and a longer trajectory (10 ns) was utilized to make the closest approximation of the binding free energy possible. [114,116,117].

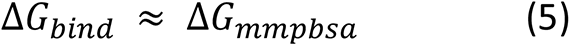

### 4.6. In silico physicochemical and ADMET profile analysis

The top lead compounds were submitted to the SwissADME webserver from the Swiss Institute of Bioinformatics to calculate their physiochemical properties and drug-likeness [118]. The lead compounds drug-likeness were evaluated based on 5 filters, the Lipinksi rule of 5, Ghose filters, Veber filter, Egan filter, and Muegge filter [119–123]. As for the ADMET analysis, admetSAR (which is also used by DrugBank to evaluate drugs ADMET profile) and ADMETlab (v2.0) webserver were collectively used to analyze each lead compound [59,89].

## 5. Supplementary Materials

S1: 98 HIV-1 WGS used in the study (.fasta), S2: MAUVE WGA of the 98 HIV-1 sequences (.fasta), S3: Alignment of the longest conserved regions within all of the 98 HIV-1 sequences (.fasta), S4: consensus sequence of the longest conserved regions within the HIV-1 genome (.fasta), S5: MM-PBSA energy term calculations (.txt), S6: SMILES of the 4 lead compounds (.txt), S7: Molecular dynamic simulation video for HIV-1 RT with Phomoarcherin B (.mp4), S8: Scorings from the additional docking studies shown in Fig 11 (.txt), and S9: The NAMD simulation setup (configuration) files used for each simulation between the HIV-1 RT-lead complex (.inp, plain-text).

## 6. Supplementary Materials

All relevant data are within the paper is provided in the paper itself and the supplementary materials.

## 7. Author Contributions

Conceptualization, N.A.G., Ö.B.; methodology, N.A.G.; software, N.A.G.; validation, N.A.G., Ö.B., B.E.S., R.S.S.; formal analysis, N.A.G.; investigation, N.A.G., Ö.B., B.E.S., R.S.S.; resources, B.E.S.; data curation, N.A.G.; writing—original draft preparation, N.A.G.; writing—review and editing, Ö.B., B.E.S., R.S.S.; visualization, N.A.G.; supervision, Ö.B., B.E.S.; project administration, Ö.B.. All authors have read and agreed to the published version of the manuscript.

## 8. Competing interests

The authors declare no competing interests.

## 9. Acknowledgments

The computational resources for performing the virtual screening were provided by the Süzek lab at Muğla Sıtkı Koçman University. The molecular dynamics simulations and MM-PBSA calculations reported in this thesis were performed on TUBITAK ULAKBIM, High Performance and Grid Computing Center (TRUBA resources). This research did not receive any monetary funding.

## Notes

### Competing Interest Statement

The authors have declared no competing interest.

